# Triple-helical ligands selectively targeting the closed αI domain of integrin α2β1

**DOI:** 10.1101/2025.07.28.667101

**Authors:** Shinsuke S. Shibata, Hiroya Oki, Kazuki Kawahara, Ryo Masuda, Kazunori K. Fujii, Takaki Koide

## Abstract

Integrins α1β1, α2β1, α10β1, and α11β1 are known to recognize the GxxGEx motif of the collagen triple helix through their αI domains, with the Glu residue engaging the domain *via* a divalent metal cation. The binding amino acid sequences containing these motifs, identified from native collagen, exhibit low subtype selectivity. Here we identify novel triple-helical peptides that selectively bind the α2I domain independently of Glu and metal ions. Yeast two-hybrid screening of randomized triple-helical peptide libraries yielded non-natural sequences in which the canonical Glu was replaced by aliphatic residues such as Met. X-ray crystal structural analysis of the representative variant GFOGMR in complex with the α2I domain revealed a new binding mode. In this mode, peptides are recognized by the closed, inactive conformation of the α2I domain without metal coordination. This cryptic interaction, likely unused by native ligands, provides a new basis for the design of subtype-specific ligands targeting collagen-binding integrins.

## Introduction

Collagen is a major protein of the extracellular matrix (ECM) of animals and is characterized by a triple-helical structure with a repeating sequence of Gly-Xaa-Yaa. Xaa and Yaa positions are often occupied by Pro and 4(*R*)-hydroxyproline (Hyp, O) residues, respectively. The latter is generated by post-translational modification of Pro, contributing to the thermal stability of the triple-helical structure (1,2). Collagen plays an important role in cell adhesion, migration, and differentiation, and these functions are driven by specific interactions between certain regions of the collagen triple helix and receptors such as integrins, as well as matricellular proteins such as osteonectin, known as secreted protein acidic and rich in cysteine (SPARC) or basement-membrane protein 40 (BM-40), and pigment epithelium-derived factor (PEDF) (3–5).

Integrins are transmembrane heterodimeric proteins composed of α and β subunits that function as cell adhesion receptors. They mediate interactions between ECM components, such as laminin and collagen, and cell adhesion molecules expressed on hematopoietic cells (6–8). Among the 18 known α subunits, nine (α1, α2, α10, α11, αD, αE, αL, αM, and αX) contain an inserted αI domain, which serves as the principal ligand-binding module (6). The αI domain is capable of reversibly adopting two distinct conformational states: a closed, low-affinity form and an open, high-affinity form. In the latter, the metal ion-dependent adhesion site (MIDAS) is exposed, facilitating ligand binding. When natural ligands (such as collagen and specific laminin isoforms) bind to the open conformation of the αI domain, the α7 helix is displaced downward, enabling association with the βI domain. This interdomain communication propagates conformational rearrangements of the extracellular domains, leading to global change of the heterodimer conformation for signal transmission across plasma membranes (Figure 1) (9).

**Figure 1.**
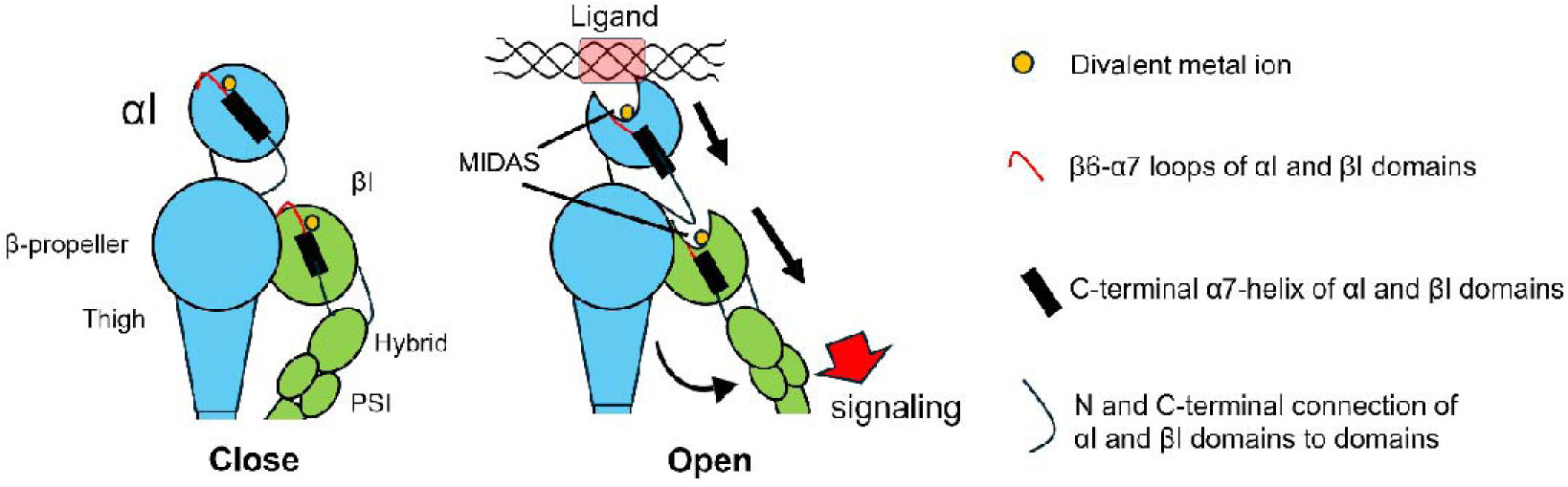
Conformational states of the extracellular headpiece of αI domain integrins. MIDAS, metal-ion dependent adhesion site; PSI, plexin-semaphorin-integrin domain.

Integrins harboring the I domain, namely α1β1, α2β1, α10β1, and α11β1, are classified as collagen receptors. Integrin α1β1 is expressed in various mesenchymal cells, and α2β1 is present on platelets and epithelial cells (10,11). Integrin α10β1 is primarily found on chondrocytes, whereas α11β1 was first identified during the differentiation of human fetal muscle cells (12,13). All the αI domains of these integrins recognize the GxxGEx (x = any amino acid) motif of the collagen triple helix *via* the divalent metal ion in the MIDAS, which coordinates to the carboxylate group of Glu (14). The integrin-binding sequences of natural collagens exhibit low selectivity for each of the four types of integrins (15,16). Therefore, identifying ligands capable of selectively binding to a particular type of integrin holds promise not only for the elucidation of each integrin’s specific functions but also for the development of integrin-targeting therapeutics.

Our research group has developed a method for obtaining triple-helical peptides capable of binding to target proteins using a combinatorial library of random peptides constructed in yeast cells and screening with a two-hybrid system (17). This library offers amino acid sequences that are more diverse than the natural triple-helical domain of collagen. In the present study, we applied this approach to the α2I domain of integrin α2β1, a receptor known to mediate platelet aggregation, with the aim of obtaining a peptide ligand with specificity for this domain. The resulting unnatural ligand exhibited specificity for the α2I domain. Furthermore, subsequent biochemical and structural analyses revealed that the specificity was attributed to a previously unexpected non-canonical mode of interaction in which the closed conformation recognized the triple helix in a metal ion-independent manner.

## Result

### Selection of integrin α2I domain-binding sequences from random triple-helical peptide library

Using the yeast two-hybrid (Y2H) system, we screened a random peptide library to identify triple-helical peptides capable of selectively binding to the integrin α2I domain. In this screening approach, the human integrin α2I domain was expressed as a GAL4 DNA-binding domain (GAL4-BD) fusion protein within yeast cells to serve as the bait. A library of GAL4 activation domain (GAL4-AD) fusion peptides was constructed as described in a previous study (17). In the library, six amino acid residues with collagen-like repeats of Gly-Xaa-Yaa were randomized (Figure 2A). Screening of 4.1 × 10^5^ clones resulted in the identification of more than 1000 binding sequences (see dataset). These amino acid sequences in selected clones were identified *via* next-generation sequencing (NGS) analysis. Figure 2A presents the frequency of amino acid residues occurring in the random region of the selected sequences. Glu, the essential amino acid residue for the binding interaction, frequently occurred at position Xaa6. Notably, aliphatic amino acids such as Met were preferred over Glu at Xaa6, and a large proportion (93%) of the individual sequences obtained had no Glu residues at Xaa positions. All sequences identified with an aliphatic amino acid at position Xaa6 were not found in human collagen α-chains.

**Figure 2.**
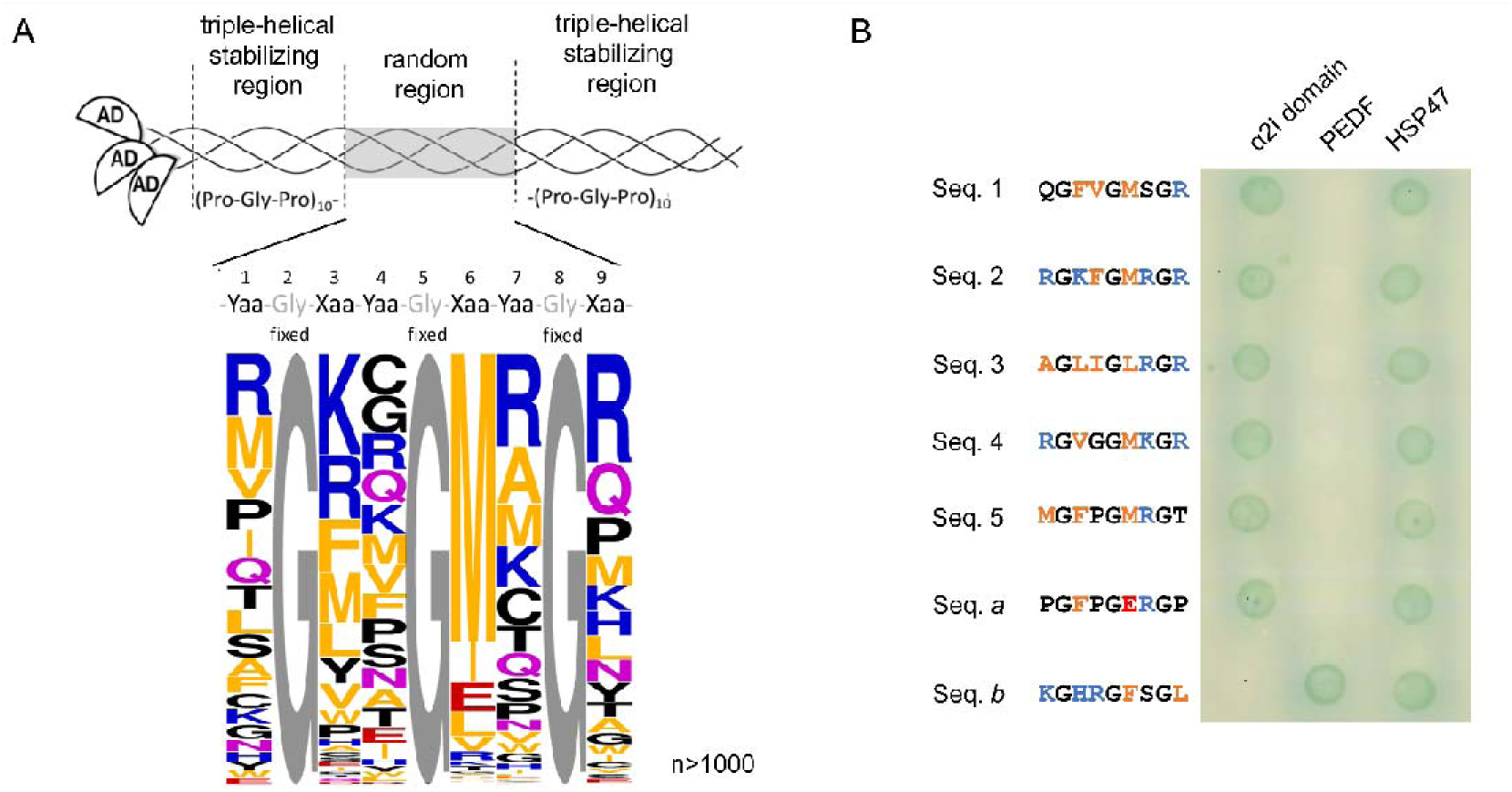
Selection of integrin α2I domain-binding triple-helical peptides. (A) Design of the library construct (upper) and Weblogo diagram showing the preferred amino acid residue at each position based on the obtained α2I domain-binding sequences from Y2H screening (lower). Gly at every 3 residues were fixed. Peptides were selected from 4.1 × 10^5^ clones at 25 °C with 200 ng/mL aureobasidin A. Sequences in hit clones were determined using NGS. The composition ratio was corrected for the theoretical occurrence rate of amino acid residues, which depended on codon combinations from the designed sequence (Table S1). In calculating the amino acid preference from the identified sequences, the occurrences of Ser, Arg, and Leu were counted with a weight of 1/3, while those of Ala, Pro, Gly, and Val were counted with a weight of 1/2, reflecting their variable positions within degenerate codons. (B) 2-hybrid interactions between AD-fusion peptides with sequences obtained and the BD-fusion α2I domain, PEDF, and HSP47. The interaction allows yeast cells to survive on the plates and turn blue by degrading X-α-Gal. Cells were cultured at 25 °C for 3 days.

Next, we evaluated the interaction between individual peptides harboring the selected sequences and the α2I domain (Figure 2B). Five representative sequences featuring aliphatic amino acids at Xaa6 were selected. Seq. 1 was the most frequently observed sequence in the selection. Seq. 2 was the sequence of a selected clone that showed the highest similarity to that composed of residues with the highest enrichment scores at each randomized position. Seq. 3 was the most frequently observed sequence among those with Leu at Xaa6. Seq. 4 contained a Gly-Gly sequence, known to significantly destabilize the triple-helical structure (Shah *et al.,* 1997). Seq. 5 contained the GFPGMR sequence, in which the Glu residue in the known integrin-binding sequence GFPGER (Seq. *a*) was replaced by Met. Here, natural collagen-derived integrin-binding sequence GFOGER was not used because proline hydroxylation does not occur within yeast cells. Seq. *b* (KGHRGFSGL), a PEDF-binding sequence, was used as a control of the Y2H interaction in combination with PEDF (5). Furthermore, heat-shock protein 47 (HSP47) was employed as an indicator of the triple-helical conformation of GAL4-AD fusion peptides. HSP47 recognizes triple-helical sequences containing Pro-Gly-Pro repeats flanking a randomized region or those with Arg at the Yaa position (19,20). All peptides containing the sequences identified through the screening engaged in two-hybrid interactions with the integrin α2I domain but not with PEDF. Additionally, all peptides exhibited interactions with HSP47, indicating that these peptides maintained a triple-helical conformation within yeast cells under the assay conditions at 25 °C. Moreover, these results strongly suggested that Glu residues at the Xaa position, which had been thought to be essential for the binding interaction with the α2I domain, could be replaced by aliphatic amino acids such as Met.

### GFOGMR-containing triple-helical peptide binds to α2I domain in a metal ion-independent manner

The results shown in Figure 2B indicated that the peptides containing the Met residue at the Xaa6 position, where only Glu was thought to be preferred, interacted with the α2I domain. To assess the relative binding affinity, we used the enzyme-linked immunosorbent assay (ELISA) to evaluate the interaction of chemically synthesized peptide polymers coated on well surfaces with recombinantly expressed glutathione *S*-transferase (GST)-tagged integrin α2I domain. The triple-helical peptide polymers were generated by polymerizing peptides containing three Cys residues at both termini *via* disulfide bonds, as previously described (21). C3-GFOGER and C3-GFOG**M**R peptides, serving as monomers for preparing the corresponding polymer, were designed to incorporate GFOGER and GFOGMR sequences, respectively, whereas C3-PRG (containing the GPOGPR sequence) was used as a negative control (Table S2). To examine the metal-ion dependency of the interaction, ELISA was performed in the presence of either MgCl_2_ or ethylenediaminetetraacetic acid (EDTA). Under MgCl_2_-supplemented conditions, the GST-fused α2I domain was bound to both polymerized poly-C3-GFOG**M**R and poly-C3-GFOGER (Figure 3A, solid lines). Notably, the interaction with polymerized poly-C3-GFOG**M**R was not inhibited by EDTA; instead, the binding affinity increased under the metal ion-depleted condition (Figure 3A, dotted lines).

**Figure 3.**
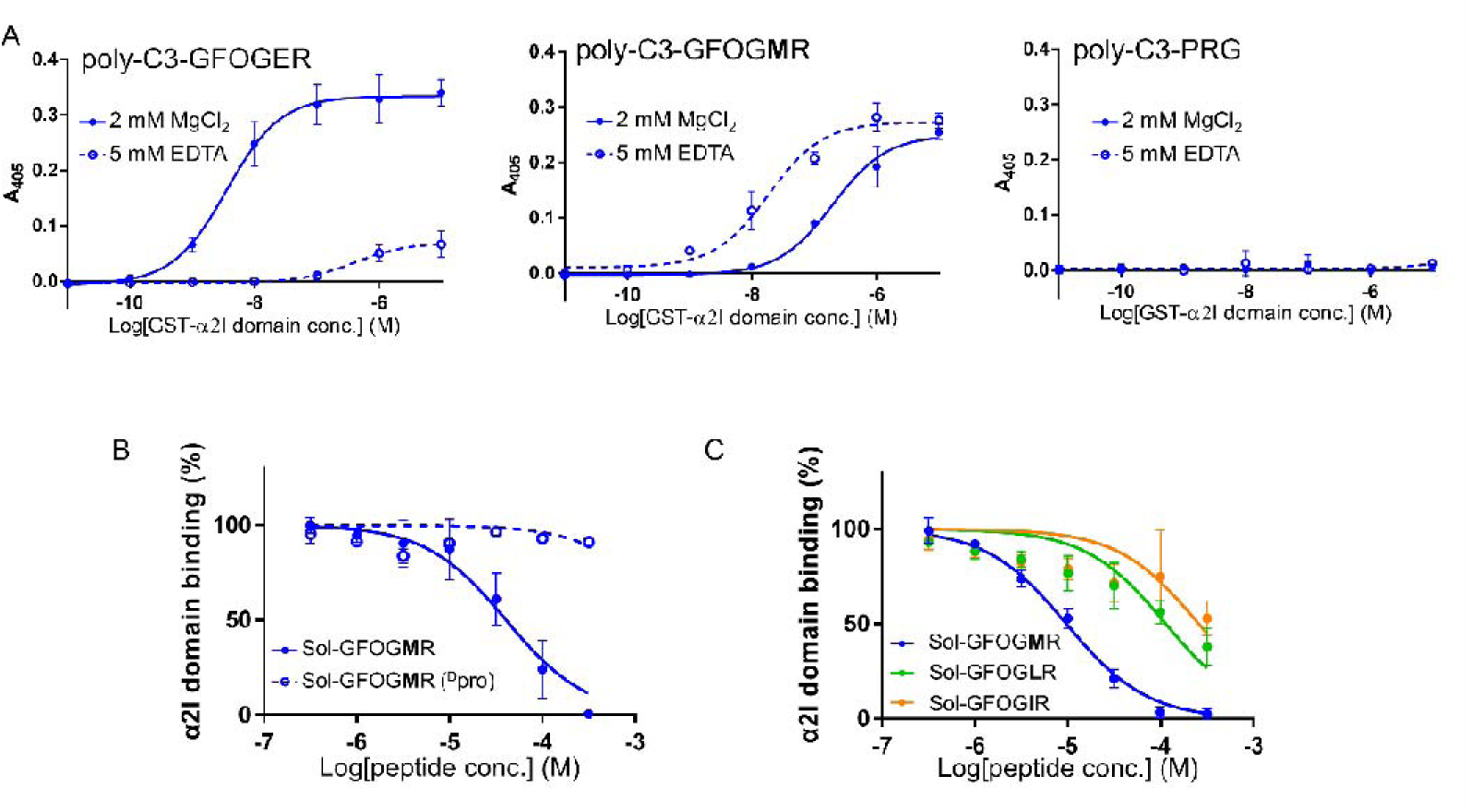
Interaction between GFOGMR-containing peptide polymer and GST-integrin α2I domain. (A) Binding of GST-integrin α2I domain to peptide polymer. Wells were coated with 10 μg/mL peptide polymer and incubated with various concentrations of GST-integrin α2I domain at 4 °C in the presence of either MgCl_2_ (solid line) or EDTA (dotted line). After incubation, an HRP-conjugated anti-GST antibody was applied, followed by staining with ABTS, and the absorbance at 405 nm was measured. (B) Investigation of the conformational tendency of the interaction. Wells were coated with poly-C3-GFOGMR (10 μg/mL) and incubated with GST-α2I domain (3 nM) in the presence of Sol-GFOGMR (solid line) or Sol-GFOGMR(DPro) (dotted line). The experiment was conducted at 4 °C in the presence of 5 mM EDTA. *n* = 3, mean ± S.D. (C) Evaluation of the affinity of the α2I domain for triple-helical peptides containing aliphatic amino acid residues. *n* = 3, mean ± S.D.

Next, we assessed the triple-helical conformation dependency of GFOGMR recognition by the GST-α2I domain using competitive ELISA with soluble peptides (Figure 3B). Peptides containing Glu interact with αI domains in a triple helical-dependent manner. Therefore, we evaluated whether the peptide containing the GFOGMR also required the triple-helical structure for the interaction. Sol-GFOG**M**R was designed as a host–guest peptide in which the GFOGMR sequence was embedded within a Pro-Hyp-Gly repeating sequence (Table S2). Additionally, Sol-GFOG**M**R(^D^Pro) was synthesized by replacing each L-proline residue within the host Pro-Hyp-Gly repeats by a D-proline (D-Pro) residue, with the intention of disturbing the triple-helical structure (Table S2). Circular dichroism (CD) spectroscopy revealed that Sol-GFOG**M**R adopted a stable triple-helical conformation at temperatures up to approximately 50 °C (Figure S3), whereas Sol-GFOG**M**R(^D^Pro) was a single chain under the same condition. Inhibition assays using these soluble peptides demonstrated that Sol-GFOG**M**R inhibited the binding of the GST-α2I domain to poly-C3-GFOG**M**R in a concentration-dependent manner, whereas Sol-GFOG**M**R(^D^Pro) had no significant inhibitory effect. This result indicated that the interaction between GFOGMR and the α2I domain was dependent on the triple-helical structure of the peptide.

Screening for α2I domain-binding sequences revealed that the Xaa6 position preferred aliphatic residues such as Ile and Leu, in addition to Met. To further investigate the impact of different aliphatic residues at this position, we evaluated the relative binding affinities of triple-helical peptides containing the GFOGxR motif (x = Leu or Ile) using similar inhibition assays (Figure 3C). The results demonstrated that the binding affinities of Sol-GFOG**L**R and Sol-GFOG**I**R were significant, although they were approximately 1/100th of the binding affinity of Sol-GFOG**M**R. This finding was consistent with the Y2H screening results (Figure 2A), in which Met was the most highly enriched residue among the aliphatic residues, suggesting that Met at the Xaa position was particularly favorable for interacting with the α2I domain.

### GFOGMR-containing triple-helical peptide is an α2I domain-selective ligand

To assess the selectivity of the triple-helical peptide containing the GFOGMR sequence for the integrin α2I domain, we evaluated its affinity for other collagen-binding integrin αI domains. The binding of four GST-fused integrin αI domains (α1, α2, α10, and α11) was assessed using peptide polymers coated onto assay wells (Figure 4). Consistent with previous reports, all four GST-integrin αI domains exhibited significant binding to poly-C3-GFOGER in the presence of MgCl_2_ (15,16). The binding affinities of α2I (blue line) and α11I (purple line) domains were highest, whereas those of α1I (red line) and α10I (green line) domains were relatively lower. Furthermore, these interactions were nullified in the presence of EDTA, confirming their metal-ion dependency. By contrast, poly-C3-GFOG**M**R exhibited significantly weaker interactions with GST-fused α1I, α10I, and α11I domains, showing only 1/100th to 1/1000th of the binding affinity observed for the GST-α2I domain, regardless of the presence of divalent metal ions. These findings establish the high selectivity of triple-helical peptides containing the GFOGMR sequence for the integrin α2I domain.

**Figure 4.**
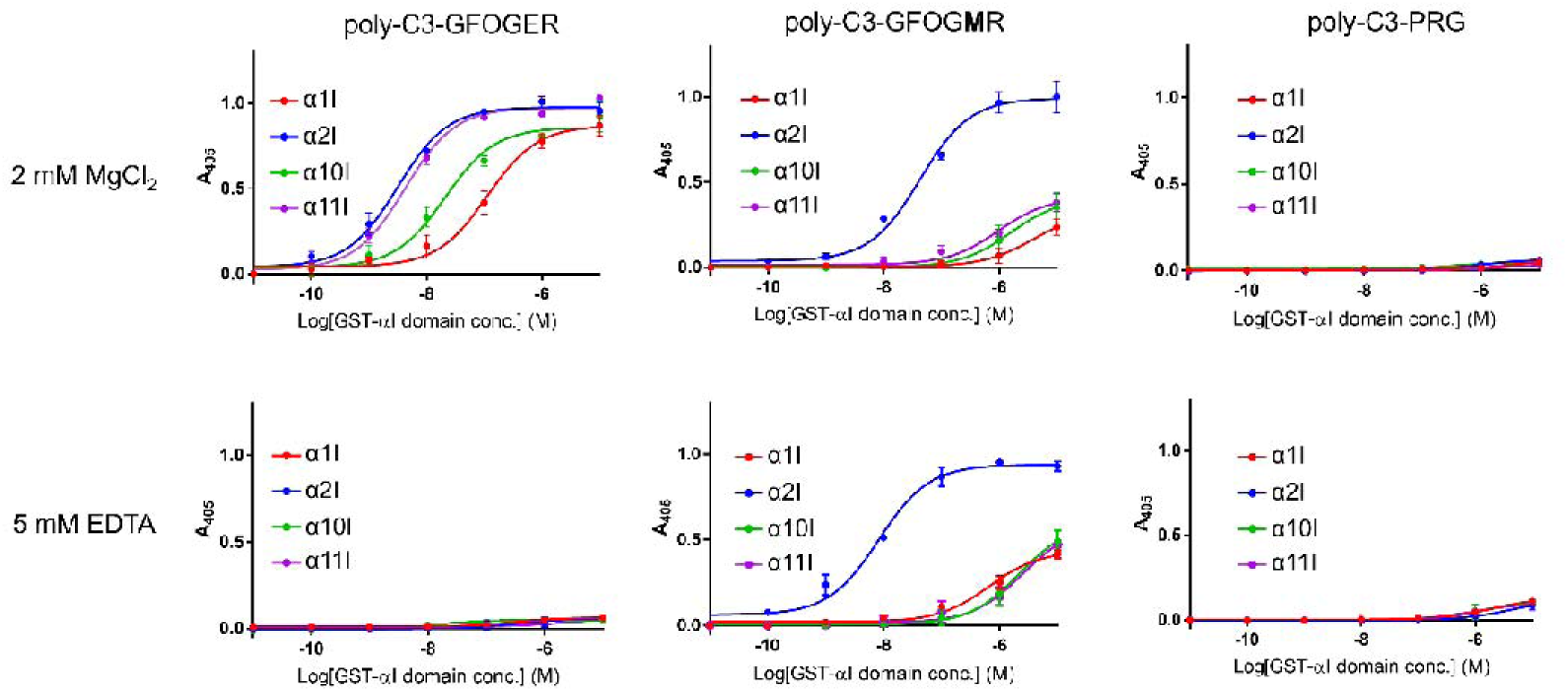
Binding of αI domains to triple-helical peptide polymer. Wells were coated with 50 μL of peptide solution (10 μg/mL). GST-integrin αI domains were incubated at 4 °C in the presence of either MgCl_2_ or EDTA. Red, blue, green, and purple lines are α1, α2, α10, and α11 I domain, respectively. *n* = 3, mean ± S.D.

### Analysis of integrin-mediated cell attachment and signal transduction on GFOGMR-coated surface

To investigate whether the triple-helical peptide containing the GFOGMR sequence could induce cell adhesion *via* integrin α2β1, we examined its ability to support the attachment of α2β1-expressing cells. HT1080 human fibrosarcoma cells express α2β1 as a major collagen receptor (22), whereas PC12 cells, a rat adrenal pheochromocytoma cell line, express α1β1 as a collagen-binding integrin but lack α2 expression (23,24). The adhesion of these cells to peptide polymers coated onto wells was assessed (Figure 5A). Both HT1080 and PC12 cells adhered to type I collagen and poly-C3-GFOGER in the presence of MgCl_2_, and this adhesion was nullified by EDTA. By contrast, HT1080 cells adhered to poly-C3-GFOG**M**R regardless of the presence of Mg^2+^, whereas PC12 cells exhibited negligible adhesion to poly-C3-GFOG**M**R. These results were consistent with those shown in Figure 4, which demonstrated that the GFOGMR-containing peptide was selectively bound to the α2I domain.

**Figure 5.**
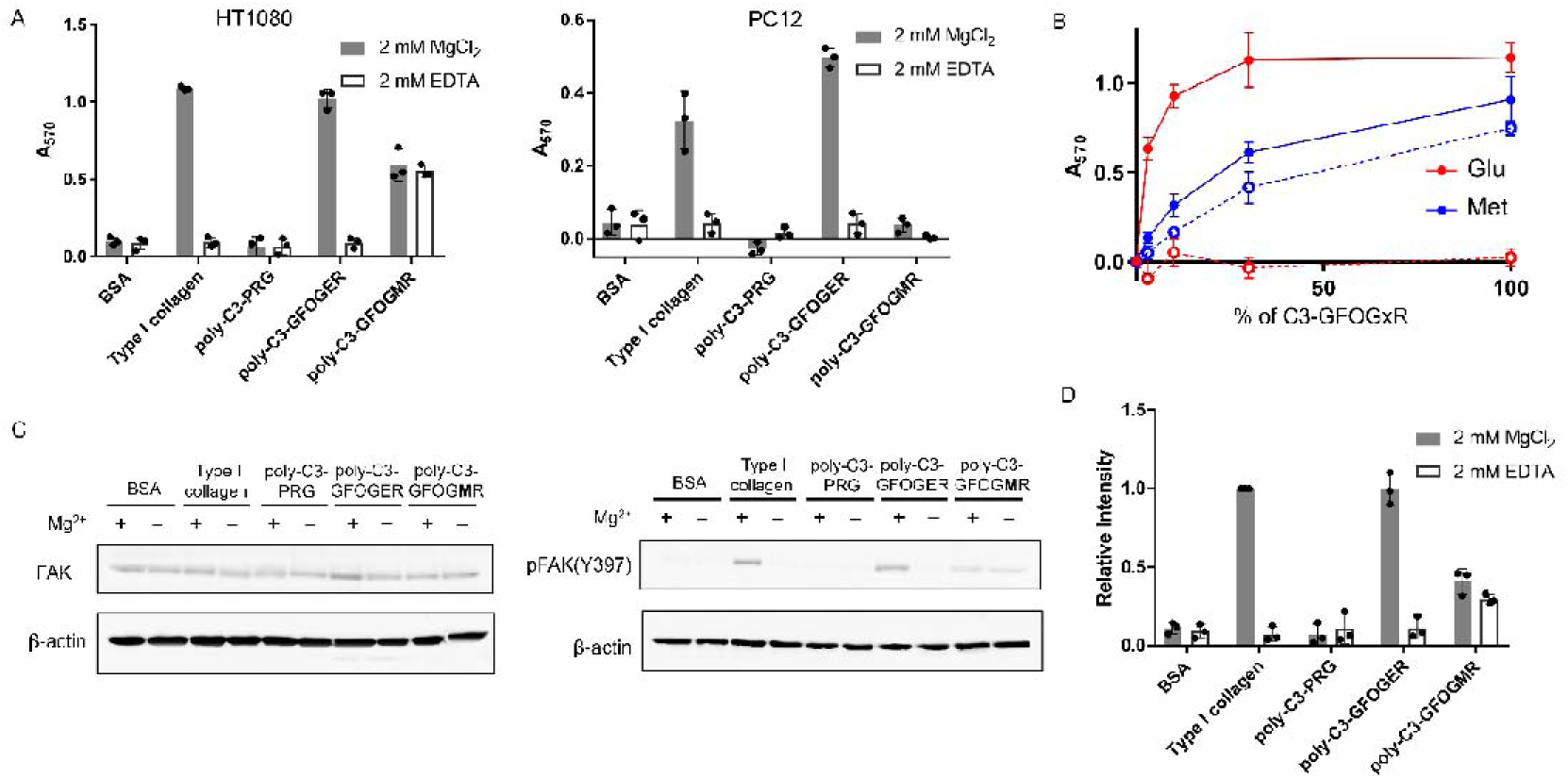
Cell adhesion to peptide polymers coated on the dishes. (A) Adhesion of HT1080 (left) and PC12 cells (right) to each peptide polymer, type I collagen, and BSA. Cells were seeded on peptide polymers, type I collagen, or BSA-coated plastic dishes (50 μg/mL solutions were used for the coating). After 2 h of incubation at each temperature, cells were immobilized with 4% *p*-formamide in phosphate buffer and stained with 5% crystal violet. *n* = 3, mean ± S.D. (B) Dependence of HT1080 cell attachment activity on the integrin α2β1-binding peptide concentration in the polymer. Percentages represent ratios of the C3-GFOGxR (x = Glu, Met) peptide weight per total peptide weight. Adhesion was obtained in the presence of either 2 mM MgCl_2_ (solid line) or 2 mM EDTA (dotted line). *n* = 3, mean ± S.D. (C) Analysis of integrin-dependent phosphorylation of FAK. HT1080 cells were seeded on peptides, type I collagen, or BSA-coated plastic dishes (50 μg/mL solutions were used for the coating). After 2 h of incubation at each temperature, cells were harvested and lysed with SDS sample buffer. Western blotting was performed with an anti-phospho-FAK (pTyr397) antibody and an anti-FAK antibody. Anti-β actin antibody blotting was used to show the amount of protein loaded on the lane. (D) Experiment of (C) was conducted three times in total, and the band intensity of phosphorylated FAK (pTyr397) relative to the band intensity of FAK under each condition was measured using ImageJ. Data from two trials, excluding Experiment A, are presented in Figure S5. Values were calculated using the band intensity of type I collagen (Mg^2+^ +) as the reference. *n* = 3, mean ± S.D.

Figure 5B illustrates the adhesion of HT1080 cells as a function of the proportion of the binding sequences within the polymers. Co-polymers comprising C3-GFOGxR (x = Glu or Met) and C3-PRG were prepared with varying C3-GFOGxR (x = Glu or Met) ratios and coated onto wells. HT1080 cell adhesion increased as the C3-GFOGxR (x = Glu or Met) content of the polymer increased. Representative images of cell adhesion under each condition are shown in Figure S4. The peptide polymer containing GFOGMR had fewer adhered cells than polymers containing GFOGER. The cell adhesion system also accommodated the metal ion-independent interaction of GFOGMR with integrin α2β1.

To determine whether HT1080 cell adhesion to poly-C3-GFOG**M**R could induce integrin-dependent cellular responses, we analyzed FAK phosphorylation by western blotting following a 2 h-incubation on peptide polymer-coated wells (Figure 5C). Type I collagen and poly-C3-GFOGER promoted FAK phosphorylation in the presence of MgCl_2_ but not in the presence of EDTA. Conversely, adhesion to poly-C3-GFOG**M**R induced relatively weak FAK phosphorylation even in the presence of EDTA. Figure 5D shows the extent of FAK phosphorylation under each condition, quantified on the basis of the intensity of phosphorylated FAK (pTyr397) relative to total FAK determined using ImageJ. Measurements were performed in triplicate, including the results shown in Figure S5. The degree of FAK phosphorylation induced by poly-C3-GFOG**M**R was markedly lower than that induced by poly-C3-GFOGER or type I collagen. These results indicated that the surface-immobilized GFOGMR-peptide acted as a weak agonist of integrin α2β1.

### X-ray crystallographic analysis reveals that the GFOGMR-containing triple-helical peptide binds to the closed α2I domain

To elucidate the structural basis of molecular recognition, we performed X-ray crystallographic analysis of the crystallized complex of a peptide containing GFOGMR (named GFOG**M**Rshort) and human integrin α2I domain in the absence of divalent metal ions. GFOG**M**Rshort consisted of a 24-residue sequence, [Ac-(GPO)_3_GFOGMR(GPO)_3_-NH_2_], and the structure of the complex was determined at a resolution of 1.6 Å (Figure S6, Table S3).

The conformation of the α2I domain of the complex remained nearly identical to that of the apo-α2I domain, particularly from the αC helix to the α7 helix (RMSD = 0.33 Å) (Figure 6A) (25). This region undergoes significant structural rearrangement upon conformational activation of the α2I domain. This result indicated that GFOG**M**Rshort was bound to the closed, and hence inactive, conformation of the α2I domain. GFOG**M**Rshort adopted a triple-helical conformation and attached to the α2I domain. This result was consistent with the competitive ELISA result shown in Figure 3B. Similar to the peptide containing GFOGER (26), GFOG**M**Rshort attached to the α2I domain by covering the MIDAS, where a divalent metal ion can be coordinated. The Met residue in the middle chain of the triple helix was inserted into the MIDAS (Figure S7).

**Figure 6.**
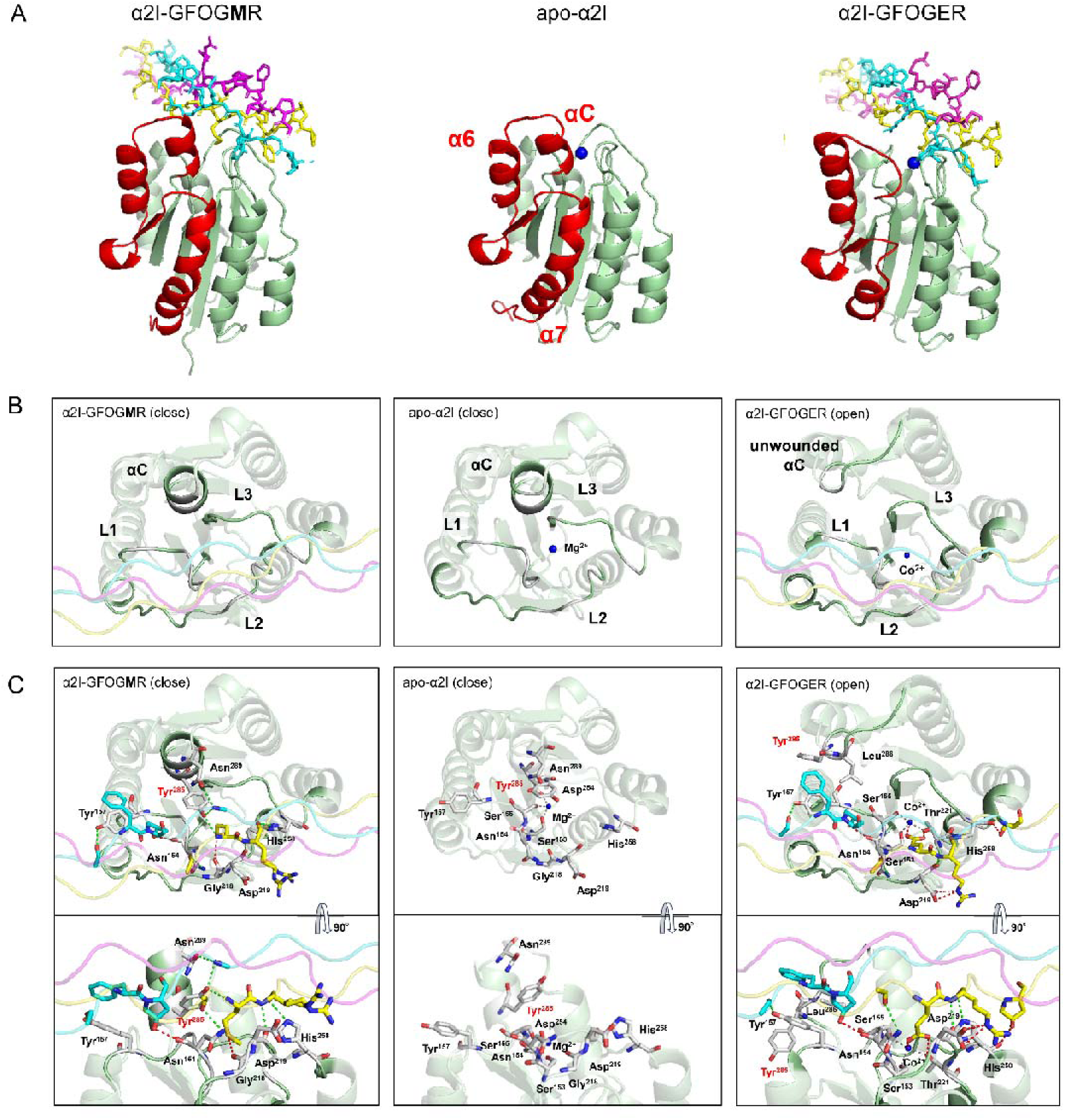
Structural determination of the α2I domain with GFOGMR-containing triple-helical peptide. (A) Comparison of the structure of the α2I domain. The complex is shown with the α2I domain as a cartoon and peptides as a stick model. Leading (L), middle (M), and trailing (T) chains in the triple helix are colored cyan, yellow, and magenta, respectively. The metal ion is shown as a blue ball. The region from the αC-helix to the α7-helix is shown in red. Crystal structure of complex with GFOGMRshort (Left), the apo-α2I domain (PDB ID code 1AOX) (Middle) and complex with the triple-helical peptide Ac-(GPO)_3_GFOGER(GPO)_2_-NH_2_ (PDB ID code 1DZI) (Right). (B) Close-up view of the binding interface. (C) Comparison of the details of the interactions. Selected side chains are shown as ball-and-stick. Interactions of the backbone of the triple-helical peptide are shown as green dotted lines, and interactions of the side chains are shown as red lines.

Figure 6B, C illustrates the detailed binding interface between the triple-helical peptide and the α2I domain. Unlike the GFOGER-containing counterpart, the GFOG**M**Rshort-containing complex had an intact αC helix in the α2I domain and no divalent metal ions at the binding site (Figure 6B). Moreover, the Met and Glu side chains of the middle chain of the triple helices occupied similar positions, and their modes of recognition were markedly different. The carboxylate anion of the Glu side chain directly coordinated with divalent metal ions, whereas the Met side chain was captured by Tyr285 of the αC helix and Gly218 of the L2 loop (Figure 6C). The side chain of Tyr285 pointed outward rather than inward from the interface upon transition to the open conformation. Comparison of additional residue interactions revealed that GFOG**M**Rshort engaged in more backbone-mediated contacts than the GFOGER-containing peptide, despite conservation of side chain-mediated interactions between the two complexes (Figure S8). Other conserved interactions were the interaction between leading-chain Phe and Tyr157, the hydrogen bond between middle-chain Hyp and Asn154, and the electrostatic interaction between middle-chain Arg and Asp219. These observations suggested that the Met(M)–Tyr285 interaction played a critical role in the recognition of the closed conformation of the α2I domain.

### Met-containing triple-helical peptides preferentially bind to the closed α2I domain

In the co-crystal analysis above, the GFOGMR-containing peptide was bound to the closed conformation of the α2I domain. To investigate whether the GFOGMR-containing peptide could also bind to the open conformation, we assessed its interaction using conformation-fixed α2I domain mutants. The E318W mutant of the α2I domain stabilizes the open conformation, whereas the T221A mutant locks the domain in the closed conformation (27,28). The binding of these GST-α2I domain mutants to peptide polymers coated on wells was evaluated using ELISA (Figure 7A). As expected, in the presence of Mg²U (solid lines), the E318W mutant (red line) exhibited a higher binding affinity for poly-C3-GFOGER than the wild-type (WT, green line). Conversely, the T221A mutant (blue line) showed a markedly reduced affinity, consistent with previous reports showing that Glu-containing triple-helical peptides preferentially bind to the open conformation of the αI domain (27,28). In contrast to WT, the T221A mutant displayed a similar binding affinity for poly-C3-GFOG**M**R both in the presence and absence of Mg^2+^ ions (dotted lines). The affinity of the E318W mutant was approximately 1/300th that of the T221A mutant. These results indicated that GFOGMR preferentially attached to the closed conformation of the α2I domain.

**Figure 7.**
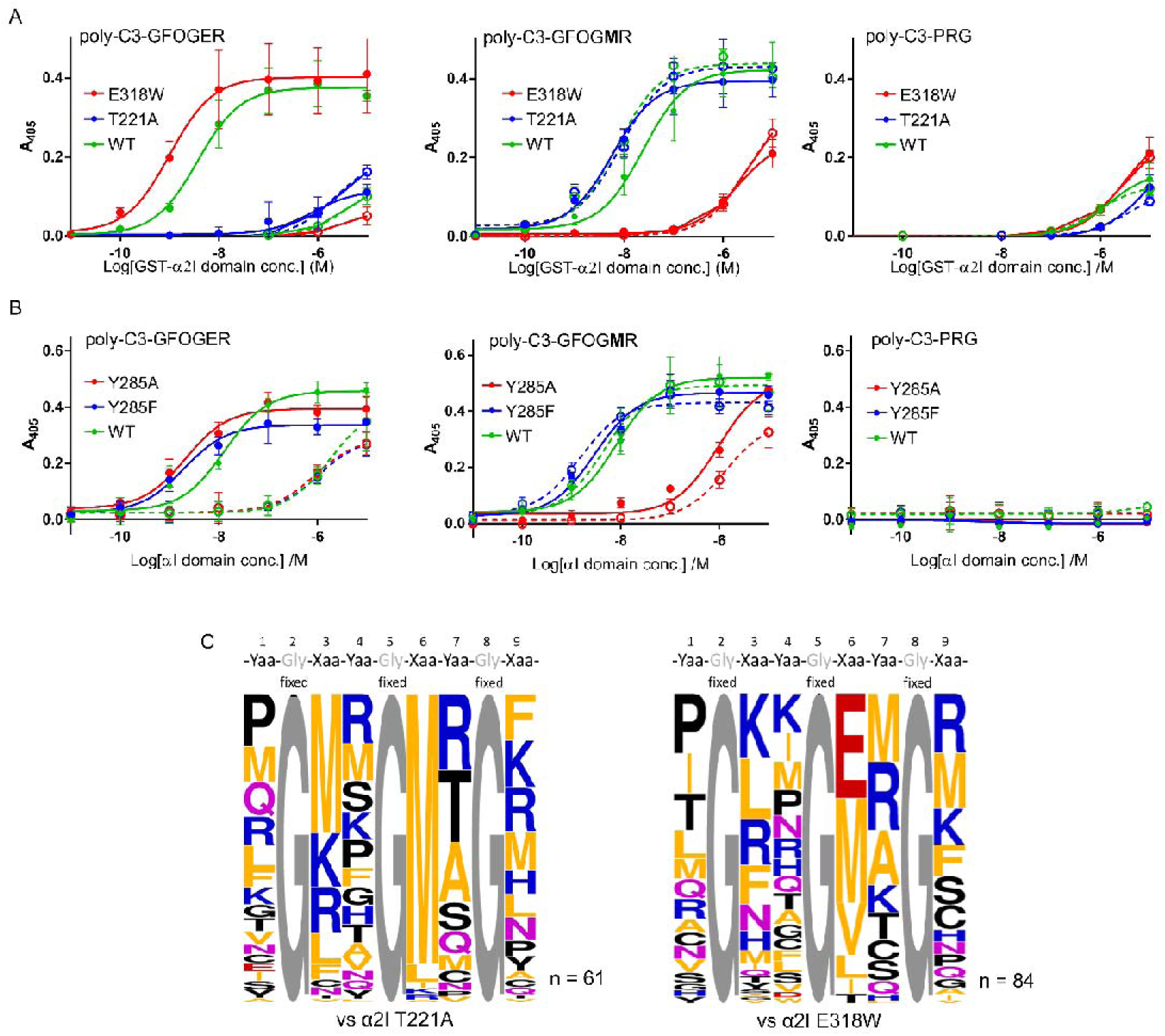
Conformational tendency of the α2I domain during interactions. (A) Interaction with surface immobilized C3-peptide polymer. Binding of constitutively open E318W (red line) and closed T221A (blue line). Adhesion was conducted in the presence of either 2 mM MgCl2 (solid line) or 5 mM EDTA (dotted line). *n* = 3, mean ± S.D. (B) Binding of α2I domain Y285 mutant. Adhesion was conducted in the presence of either 2 mM MgCl_2_ (solid line) or 5 mM EDTA (dotted line). *n* = 3, mean ± S.D. (C) Weblogo diagram showing the preferred amino acid residue at each position based on α2I domain mutant-binding amino acid sequences obtained from Y2H screening. For the T221A mutant, sequences were selected from 5.1 × 10^5^ clones at 25 °C with 200 ng/mL aureobasidin A. For the E318W mutant, sequences were selected from 2.7 × 10^5^ clones at 25 °C with 200 ng/mL aureobasidin A. Among the hit clones, 96 clones were analyzed using the Sanger method. The composition was corrected for the occurrence rate of amino acid residues, which depended on codon combinations from the selected sequence.

Furthermore, the co-crystal structure suggested that the Met side chain of the peptide interacts with Tyr285 on the αC-helix, which unravels upon transition to the open conformation. To assess the importance of Tyr285, mutants in which this residue was substituted with Phe or Ala were constructed, and their binding to the peptide polymers coated on wells was evaluated (Figure 7B). The Y285F mutant showed little change in affinity for poly-C3-GFOG**M**R, whereas the Y285A mutant showed a marked reduction compared with WT. In contrast, both mutants bound poly-C3-GFOGER with affinities similar to WT. These results demonstrate that Tyr285 is critical for binding of the GFOGMR peptide and provide evidence supporting the preferential recognition by the closed conformation of the α2I domain.

On the basis of these findings, we revisited a Y2H screening for binding sequences targeting each α2I domain mutant stabilizing the open or closed conformation. Screening a library of 5.1 × 10^5^ diverse clones against the α2I T221A mutant yielded over 4000 positive clones. By contrast, screening a library of 3.5 × 10^5^ clones against the α2I E318W mutant produced approximately 1000 positive clones. Sequence analysis of 92 positive clones for each mutant identified 61 binding sequences for the α2I T221A mutant and 69 binding sequences for the α2I E318W mutant (Tables S4, S5). Figure 7C presents the preference of amino acid residues occurring in the random region of the selected sequences. Notably, 90% of the sequences obtained for the α2I T221A mutant contained either Met or Leu at position Xaa6. All other identified sequences carried Met at Xaa3, and no sequences harbored Glu at any Xaa position. By contrast, for the α2I E318W mutant, Glu was the most strongly enriched residue at Xaa6. Although aliphatic residues also enriched at this position, their enrichment was markedly lower than that observed in WT-targeted screening (Figure 2A). This distribution aligned with the difference in binding preferences between the α2I mutants, as shown in Figure 7A. Therefore, the results obtained from WT-targeted screening (Figure 2A) likely represented a merged outcome of screenings targeting both the open and closed conformations of the α2I domain.

## Discussion

In this study, we identified triple-helical ligands capable of selectively binding to the integrin α2I domain. These novel ligands substitute the Glu residue—previously considered essential for metal ion-dependent recognition—with the Met residue, thereby challenging the prevailing model of the integrin’s ligand recognition mechanisms. Conventional strategies for identifying integrin-binding sequences have focused on native collagen as a target. Hook and colleagues mapped α2β1 and α1β1 integrin-binding sites on human type I and III collagens using electron microscopy (29,30). Farndale’s group developed synthetic triple-helical peptide libraries (named Collagen Toolkits) to cover the entire sequences of type II and III collagen α-chains and identified binding sequences for integrin α1I, α2I, and α10I domains (31–34). All sequences identified through these studies contain the GxxGEx motif and are characterized by low subtype specificity. These studies establish a structural framework in which Glu plays a central role in the metal-dependent recognition of the collagen triple helix by integrins.

Conversely, we employed a combinatorial screening strategy using a randomized triple-helical peptide library, enabling access to a broader chemical space unrestricted by the sequence context of native collagens (17). This approach yielded peptide sequences containing Met—an aliphatic residue—at the Xaa position, which selectively attached to the α2I domain. Met is markedly less common at the Xaa position of native collagen sequences than Glu, which is canonical in the integrin-binding sequences of the triple-helical regions of collagen. In fact, within the triple-helical regions of human fibrillar collagen α-chains [α1(I), α2(I), α1(II), α1(III), α1(V), α2(V), α3(V), α1(XI), and α2(XI)], GEx motifs appear 385 times (13.3%), whereas GMx motifs are found only 24 times (0.83%) (17,35). In this study, no sequences containing GxxGMx motifs identified by screening against the WT α2I domain and its mutants were found in human collagen sequences. Furthermore, structural analyses revealed that these Met-containing motifs were preferentially recognized by the α2I domain in its closed, signaling-inactive conformation. These findings suggest that such a Met-mediated binding mode is unlikely engaged in the endogenous ECM-mediated signaling system and may instead represent a cryptic recognition of the integrin.

To further elucidate the interaction between Met-containing triple-helical peptides and the α2I domain, we conducted biochemical and structural analyses using a Met-substituted triple-helical peptide (GFOGMR), in which the canonical Glu residue of GFOGER was replaced by Met. This peptide attached to α2I *via* a novel recognition mode that did not require the previously essential divalent metal ion (Figure 3A). Subsequent X-ray crystallographic analysis revealed the structural basis of this alternative interaction. GFOGMR-containing triple-helical peptide interacted with the α2I domain in its closed conformation, which had been presumed to lack triple-helix recognition and signaling capability (Figure 6). In the complex of the peptide with the α2I domain, the Met residue in the middle strand of the GFOGMR-containing peptide is positioned adjacent to the MIDAS, similar to the positioning of Glu in GFOGER; however, the mode of interaction is distinct. Whereas the Glu carboxylate directly coordinates the metal ion (Figure 6C and Figure S9), the amide group of the main chain Met forms a hydrogen bond with the hydroxyl group of Tyr285 in the αC helix, and the side chain engages in hydrophobic interactions with the aromatic ring of Tyr285. This interaction occurs exclusively in the closed conformation of the α2I domain and underlies the peptide’s conformational selectivity, as the Tyr285 is relocated from the binding interface to an outward orientation upon the conformational change. In addition to the interaction with Tyr285, the sulfur atom of Met is positioned close to the carbonyl group of Gly218 in the L2 loop, consistent with the formation of a twisted chalcogen bond (Figure S9) (36,37). This interaction may contribute to the preferential binding of the domain to Met among the aliphatic amino acids.

Binding to the closed α2I conformation appears to rely more heavily on peptide backbone-mediated interactions than on side-chain contacts (Figure S8). This shift likely relaxes the sequence constraints of integrin ligands and permits broader sequence diversity. In support of this idea, screening against the WT α2I domain yielded a higher number of Met-containing sequences than Glu-containing sequences (Figure 2A). In addition, screening against the T221A mutant, which favors the closed conformation, yielded approximately 4000 positive clones—fourfold more than the approximate 1000 clones recovered by screening against the E318W mutant, which mimics the open conformation (Figure 7C).

The triple-helical peptide containing GFOGER interacts with the open αI domains in a metal-ion dependent manner *via* the MIDAS. This peptide shows low subtype selectivity. In contrast, the corresponding Met-containing peptide, GFOGMR peptide, is a selective ligand for the α2I domain (Figure 4). This selectivity arose from preferential recognition of the closed α2I domain. Figure 8 shows the sequence comparison among the αI domains. Tyr285 in the α2I domain corresponds to Ser284 in the α1I, His284 in the α10I, and Tyr301 in the α11I domain. Since Tyr285 in the α2I domain is critical for the interaction with the GFOGMR peptide (Figure 7B), the absence of the Tyr in the α1I and the α10I domains accounts for their markedly reduced affinities. In contrast, the Tyr residue is conserved in the a11I domain, suggesting that the low affinity of the peptide for the a11I domain may arise from other structural features. One possible reason is steric hindrance restricts the access of the triple helix to the binding site. Asn289, which is located proximal to Tyr285 in the α2I domain, corresponds to a bulky Arg305 in the α11I domain (Figure S10B). However, the a2I mutant (N289R) showed little reduction in affinity for the peptide compared to the WT (Figure S10C). This result did not support the hypothesis, leaving the structural basis for the origin of selectivity for the a11I domain unclear.

**Figure 8.**
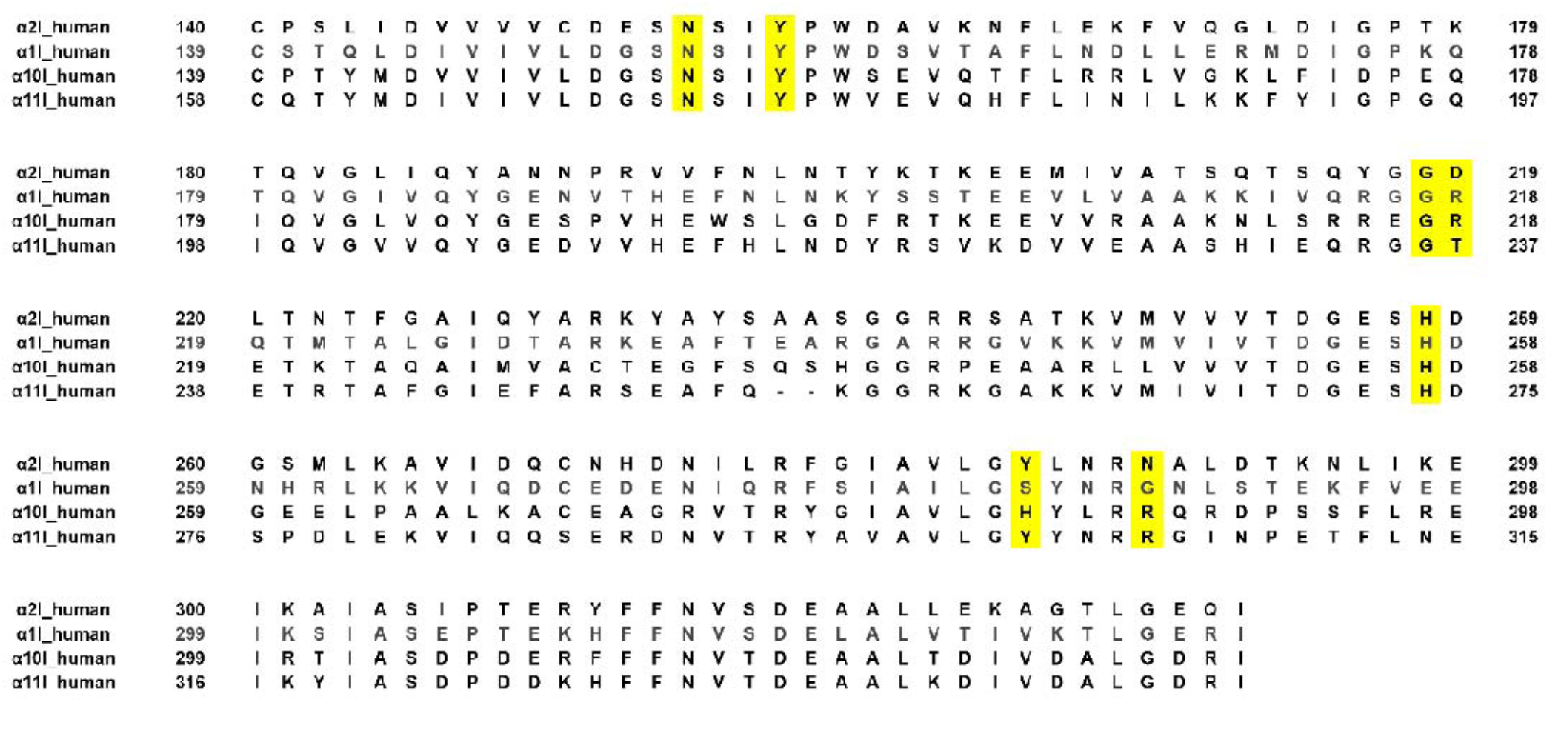
Comparison of collagen receptor integrin αI domains. Residues of the α2I domain interacting with the triple-helical peptide containing GFOGMR, as well as corresponding residues in other αI domains, are highlighted in yellow.

Despite its preferential binding to the closed conformation of the α2I domain, the triple-helical peptide containing GFOGMR was found to function as a weak agonist in cellular assays, as evidenced by its significant but weak induction of FAK phosphorylation in HT1080 cells (Figure 5C, 5D). Given that integrin signaling requires the αI domain to transition to an open conformation and cooperate with the activation of the β subunit, this observation appears paradoxical. However, this peptide retains the capacity to bind, albeit with low affinity, to the open conformation of α2I, which likely accounts for its signaling activity. Consistent with this observation, this peptide also binds to the open conformation-mimicking E318W mutant, with an affinity much lower than its binding to the closed conformation-stabilizing T221A mutant (Figure 7A). Furthermore, in the screening against the E318W mutant, Met-containing sequences emerged as the second most frequent class following canonical Glu-containing sequences, supporting this interpretation (Figure 7C).

Integrin α2β1 plays a crucial role in platelet adhesion to collagen under flow and in platelet aggregation, acting upstream of GPVI-mediated activation (38,39). Overexpression of α2β1 promotes pathological thrombosis, whereas its inhibition reduces platelet adhesion without impairing platelet activation (40). Ligands that selectively bind to closed αI domains are expected to act as inhibitors of integrins lacking agonistic activity (41). Thus, the Met-containing α2I domain-selective ligand may serve as a foundation for the development of novel antithrombotic agents, particularly through optimization aimed at enhancing affinity while minimizing the remaining agonist activity. Because the Met-containing ligand selectively binds to the α2I domain through its ability to target the closed form, ligands that selectively bind to other αI domains may also be obtained by targeting the closed forms of these domains. The ability to isolate ligands that are capable of selectively binding to a particular type of collagen-binding integrin offers not only opportunities for the development of selective inhibitors but also a powerful means to dissect the distinct physiological roles of these receptors, which have remained largely elusive. The combinatorial screening strategy employed here—based on a triple-helical peptide library and yeast two-hybrid selection—provides a valuable approach toward this goal.

## Materials & Methods

### DNA construction

Plasmids encoding human integrin α1, α2, α10, or α11 I domain were synthesized by FASMAC (Kanagawa, Japan). These constructs were inserted into the pUCFa vector with *EcoR*I and *Sal*I sites, containing α1I (Pro129-Ile334), α2I (Cys140-Gly337), α10I (Pro140-Gly336), or α11I (Cys158-Gly353) (16, 42–44). DNA fragments encoding the αI domains and a pGEX5x-1 vector were ligated after digestion with *EcoR*I and *Sal*I, and transformed into HST08 competent cells.

To prepare a plasmid encoding GAL4-BD fusion human integrin α2I domain in pGBKT7, DNA fragment encoding the α2I domain and a pGBKT7 vector were ligated after digestion with *EcoR*I and *Sal*I, followed by transformation into HST08 competent cells. To prepare plasmids encoding α2I domain mutants, inverse PCR was performed with the above GST-fused α2I domain in a pGEX-5x-1 vector as a template by Primestar MAX (TaKaRa, Shiga, Japan). The PCR product was ligated with Gibson assembly reagents (New England Biolabs, Hertfordshire, UK), followed by transformation into HST08 competent cells. Used primers are listed in Table S6. For crystallization, the α2I domain was mutated at the *N*-terminal, replacing Cys140 by Ser140, and vector-derived residues (GIPEF) remaining at the *N*-terminus were removed using inverse PCR with primers α2I cryst N forward and α2I cryst N reverse, as shown in Table S7. Subsequently, to remove vector-derived residues (VDSSGRIVTD) at the *C-*terminus, inverse PCR was performed using primers α2I cryst C forward and α2I cryst C reverse, introducing a stop codon immediately after the *C*-terminal Gly337 of the integrin α2I domain.

### Y2H screening

Y187 and Y2H gold cells in the yeast two-hybrid kit (TaKaRa) were used as host cells expressing GAL4-AD-fusion triple-helical peptides and BD-fusion integrin α2I domain, respectively. Transformation of plasmids into yeast cells was performed according to the manufacturer’s protocol. Screening of the integrin α2I domain-binding peptides was performed according to a reported protocol (17). Briefly, the Y2H gold strain expressing the integrin α2I domain was cultured to 0.5–0.6 OD_600_ in SD/-Trp broth at 30 °C. This culture was mixed with Y187 expressing AD-fusion triple-helical peptides with random sequences and filtrated through a nylon membrane filter (0.45 μm pore; GE Healthcare Life Sciences, Pittsburgh, PA, USA). After incubation of the membrane on a yeast peptone dextrose adenine (YPDA) plate at 30 °C overnight, cells were collected with saline and inoculated on quadruple-dropout (QDO; SD/-Leu-Trp-His-Ade) plates with 200 ng/mL aureobasidin A. Evaluation of interactions with individual sequences was performed according to the reported method after ligation of the ssDNA (Table S8) to the pGADT7-AD vector. For Seq. *b*, we applied a previously reported protocol (17).

### DNA sequencing

To identify the sequence of random regions using NGS, all colonies growing on QDO plates were collected into phosphate-buffered saline (PBS). The collected yeast cells were washed with PBS three times and suspended in yeast lysis buffer [50 mM Tris-HCl (pH 8.0), 150 mM NaCl, 5 mM EDTA, 1% NP-40, and 1% SDS]. The suspension was mixed with the same volume of phenol/chloroform/isoamyl alcohol (25:24:1, pH 7.9) and vortexed with glass beads for 1 min. DNA was purified from the upper layer by ethanol precipitation. Amplification of PCR products for amplicon sequencing and analysis using NGS were performed according to a reported protocol (17).

To identify the sequence of random regions using the Sanger method, colonies growing on QDO were individually collected, and PCR was performed using the primers listed in Table S9. PCR products were purified using MagExtractor–PCR & Gel Cleanup (TOYOBO, Tokyo, Japan). Amplification and analysis of PCR products for sequencing were performed by FASMAC (Kanagawa, Japan).

### Peptide synthesis

C3-GFOGER and C3-PRG were used as previously reported (21). Peptides were manually synthesized according to the 9-fluorenylmethoxycarbonyl (Fmoc)-based solid-phase method. C3-GFOG**M**R was constructed on 2-chlorotrityl chloride resins (Peptide Institute, Osaka, Japan). Sol-peptides were constructed on Rink-amide resins (Novabiochem, San Diego, CA, USA ). The coupling reaction was performed at room temperature for 1.5 h with 3–5 equivalents of Fmoc-amino acid and 1-hydroxybenzotriazole (HOBt), *N,N*′-diisopropylcarbodiimide (DIC) in *N,N*-dimethylformamide (DMF). Fmoc deprotection was performed with 20% (v/v) piperidine in DMF for 20 min. The protected peptide resin was treated with trifluoroacetic acid (TFA)/H_2_O/*m*-cresol/thioanisole/ethanedithiol (82.5/5/5/5/2.5, v/v) for 4 h at room temperature. All peptides were purified by RP-HPLC using a Cosmosil 5C18 AR-II column (20 mm id × 250 mm, Nacalai Tesque, Kyoto, Japan) and characterized using matrix-assisted laser desorption-ionization mass spectrometry (Autoflex MAX, Bruker, Billerica, MA, USA) or electrospray ionization time-of-flight mass spectrometry [Compact (ESI), Bruker] (Figure S2). All measured masses agreed with the theoretical values. The concentration of sol-peptides was measured from the absorbance of Tyr residues at 280 nm (47). C3-peptides were polymerized as described (Ichise *et al*., 2019). Peptides were annealed in 0.05% TFA/H_2_O. To establish disulfide cross-links, dimethyl sulfoxide (DMSO) was added to the peptide solution up to a final concentration of 10% (v/v), and the solution was incubated at 4 °C for a week.

### CD spectrometry of synthesized peptides

CD spectra were recorded on a J-820 CD spectropolarimeter (JASCO, Tokyo, Japan) equipped with a Peltier thermal controller using quartz cuvettes (0.5 mm path length) and connected to a data station for signal averaging. Peptides in H_2_O were heated at 95 °C for 5 min, incubated at room temperature for 10 min, and stored at 4 °C overnight to allow them to fold. Data was obtained using continuous wavelength scans from 190 to 260 nm (Figure S4). Spectra are reported in terms of ellipticity units per mole of amino acid residue [θ]_mrw_. The triple-helix melting curve was monitored by observing each peptide [θ]_225_ value while increasing the temperature from 4 to 85 °C at 18 °C h^–1^.

### GST-integrin I domain expression and purification

Clones of BL21(DE) expressing each GST-αI domain were incubated in lysogeny broth (LB) containing 100 μg/mL ampicillin at 37 °C with shaking. After the incubated solution reached the absorbance (at 600 nm) of 0.7–1.2, 0.1 mM isopropyl-1-thio-*β*-_D_-galactopyranoside (IPTG) was added, and the solution was incubated at 25 °C for 4 h. The incubated solution was centrifuged (1660 × *g*) at 4 °C for 10 min to discard the supernatant. The obtained pellet was treated with PBS and centrifuged under the above condition (4 °C, 1660 × *g*, 10 min). The cells were lysed in a lysis buffer [1 mg/mL lysozyme, 50 mM Tris-HCl (pH 8.0), 5 mM EDTA, 150 mM NaCl, 30% (v/v) glycerol]. Protease inhibitors and 0.15% NP-40 were subsequently added, and the mixture was rotated for 15 min. After mixing genomic DNA by sonication and centrifugation, the lysate was loaded onto GST-accept (Nacalai Tesque) for 1 h. After discarding the supernatant, the pellet was washed with PBS, 0.4 M NaCl in PBS, and PBS. Then, the pellet was loaded into the column tube.

To obtain the GST-integrin aI domain, fraction tubes were divided by adding GSH buffer [50 mM Tris-HCl (pH 8.0), 10 mM GSH]. After dialysis with tris-buffered saline (TBS), the protein solution was frozen with liquid N_2_ and stored at –24 °C until use. The concentration of the proteins was estimated using the Bradford assay. Bovine serum albumin (BSA) was used as the standard. The purity of GST-aI domains was checked using 12% SDS-PAGE (Figure S11).

### ELISA

Wells of 96-well plates (161093, Nunc, Kamstrup, Denmark) were coated with 50 μL of 10 μg/mL peptide polymers at room temperature for 1 h. After washing with the ELISA buffer (1 mg/mL BSA in TBS), wells were blocked with 100 μL of 20 mg/mL BSA in TBS at room temperature for 1 h. The wells were washed with the ELISA buffer three times, followed by the addition of 50 μL of 2 mM MgCl_2_ or 5 mM EDTA and the ELISA buffer containing the recombinant GST-integrin αI domain. The plate was incubated at 4 °C for 90 min and then washed with the ELISA buffer five times. Horseradish peroxidase (HRP)-conjugated anti-GST antibody (diluted 1:3000; GE Healthcare Life Sciences) was then added, followed by incubation at 4 °C for 45 min. After washing the wells three times, 50 μL of 0.5 mg/mL 2,2′-azinobis(3-ethylbenzthiazoline-6-sulfonic acid) diammonium salt (Fujifilm Wako Pure Chemical Industries, Osaka, Japan) in citrate-phosphate buffer [0.1 M citrate, 0.2 M Na_2_HPO_4_ (pH 5.0)] was added to each well. Plates were incubated at 37 °C for 20 min, and the absorbance at 405 nm was measured.

In the competitive assay, the final concentration of the GST-integrin α2I domain was 3 nM. Sol-peptides and GST-integrin α2I domain were diluted with the ELISA buffer.

### Cell culture

Cells were cultured in Dulbecco’s modified Eagle’s medium (DMEM) containing 10% fetal bovine serum (FBS; Thermo Fisher Scientific, Waltham, MA, USA), 100 units/mL penicillin, and 100 μg/mL streptomycin (Sigma–Aldrich). Cells were maintained at 37 °C in a humidified 5% CO_2_/air atmosphere. DMEM (low glucose; Fujifilm Wako Pure Chemical) was used for HT1080, while DMEM/Ham’s F12 (Fujifilm Wako Pure Chemical) was used for PC12.

### Cell adhesion assay

Cells were detached from dishes with 2 mM EDTA/PBS and recovered by centrifugation. After washing with PBS, HT1080 cells were resuspended at a concentration of 2 × 10^5^ cells/mL in PBS supplemented with either 2 mM MgCl_2_ or 2 mM EDTA. Wells of 96-well plates (Nunc) were each coated with 50 μL of 50 μg/mL synthetic peptide solution or acid-soluble type I collagen derived from bovine dermis (Koken Co., Ltd., Tokyo, Japan) overnight at room temperature. After blocking with 100 μL of 5 mg/mL BSA, the wells were washed with PBS. The cell suspension (100 μL) was then added to each well and incubated for 1 h at 37 °C. PC12 cells were cultured for 2 h at a concentration of 5 × 10^5^ cells/mL. Adherent cells were stained with 2% crystal violet in methanol for 15 min. After washing, elution was performed with 100 μL of 50% MeOH/H_2_O, and the absorbance at 570 nm was measured.

### Western blotting

After HT1080 cells (2.0 × 10^5^ cells in 2 mM MgCl_2_/PBS or 2 mM EDTA/PBS) were incubated in coated 6-well plates (Nunc) at 37 °C, they were harvested and lysed in 150 μL of the sample buffer (50 mM Tris–HCl, pH 6.7, 10% glycerol, 2% SDS). Protein concentrations were measured using a Pierce BCA protein assay kit (Thermo Fisher Scientific). After separation by SDS-PAGE on 10% gels, proteins were electrophoretically transferred to nitrocellulose membranes. The membranes were blocked with 5% non-fat milk in TBS and incubated with either 1:2000 mouse monoclonal antibody against focal adhesion kinase (FAK; clone 4.47, Merck Millipore, Billerica, MA, USA), 1:400 rabbit polyclonal antibody against phospho-FAK (pTyr397) (Invitrogen, Carlsbad, CA, USA), or 1:2000 mouse monoclonal antibody against β-actin (Sigma–Aldrich, St. Louis, MO, USA) for 1 h. HRP-conjugated anti-rabbit IgG (1:800; Promega, Fitchburg, WI, USA) or anti-mouse IgG antibodies (1:3000; Santa Cruz Biotechnology, Dallas, TX, USA) were used as a secondary antibody and applied for 30 min. Protein bands were detected using Pierce ECL Plus Western blotting substrate (Thermo Fisher Scientific) and visualized using a LAS-3000 luminescence image analyzer (Fujifilm corp., Tokyo, Japan).

### Crystallization

BL21(DE) expressing crystallization-grade α2I domain was cultured in LB medium containing 100 μg/mL ampicillin at 37 °C until the OD_660_ reached 0.6. IPTG was then added to a final concentration of 0.2 mM, and protein expression was induced overnight at 20 °C. After cultivation, the bacterial culture was centrifuged (4,000 × *g*) for 10 min at 4 °C to obtain a cell pellet. The obtained cells were resuspended in lysis buffer [20 mM Tris-HCl (pH 8.0), 1 M NaCl,], followed by the addition of lysozyme (final concentration: 0.2 mg/mL) and incubation with stirring for 15 min. Triton X-100 was then added to a final concentration of 0.2% (v/v), and the mixture was further stirred for 15 min. The cells were disrupted by sonication, and the lysate was centrifuged (24,000 × *g*) for 1 h at 4 °C to collect the supernatant. The supernatant was applied to a GSTrap HP column pre-equilibrated with the lysis buffer. The column was washed with the lysis buffer and wash buffer [20 mM Tris-HCl (pH 8.0), 150 mM NaCl]. GST-tagged α2I domain was then eluted using an elution buffer [20 mM Tris-HCl (pH 8.0), 150 mM NaCl, 20 mM GSH]. The collected protein solution was subjected to buffer exchange by dialysis against a dialysis buffer [20 mM Tris-HCl (pH 8.0), 150 mM NaCl, 5 mM CaCl_2_] and treated with Factor Xa. The enzyme-treated sample was applied to a GSTrap column again, and the flow-through fraction was collected. Further purification was performed using a Superdex 75 (16/600) gel filtration column equilibrated with an equilibration buffer [20 mM Tris-HCl (pH 8.0), 150 mM NaCl, 2 mM EDTA] to obtain the integrin α2I domain. The purified α2I domain was mixed with annealed GFOG**M**Rshort at a molar ratio of 1:1.5, and the mixture was applied to the Superdex 75 (16/600) column to collect the fractions containing the peak of the α2I domain-GFOG**M**Rshort complex. The obtained complex solution was concentrated to 10 mg/mL and subjected to crystallization screening using the sitting-drop vapor diffusion method at 4 °C under 96 conditions from Crystal Screen 1 & 2 (Hampton Research). Initial crystals were obtained under condition 22 (0.2 M ammonium acetate, 0.1 M sodium acetate buffer pH 4.6, 30% PEG 4000).

Optimization of the crystallization conditions resulted in diffraction-quality crystals from 2 µL of drops consisting of an equal mixture of the complex solution and reservoir solution (0.2 M ammonium acetate, 0.1 M sodium acetate buffer pH 4.6, 30% PEG 4000).

### Data collection and structure determination and refinement

X-ray diffraction experiments were conducted at the BL45XU beamline of the large-scale synchrotron radiation facility, SPring-8, using a PILATUS 6M detector. Diffraction data were processed using the XDS program (46). The initial phase was calculated using the molecular replacement method with the PHASER program of PHENIX (47). As the search model, we used the complex structure of the integrin α2I domain with a collagen sequence containing the WT binding motif (PDB ID code: 1DZI)(26). The analysis resulted in an initial model structure containing four molecules of the integrin α2I domain-GFOG**M**Rshort complex per asymmetric unit. Subsequent iterative structure refinement was performed using phenix.refine and the Coot program, ultimately yielding a final complex structure with *R_work_* and *R_free_* values of 0.192 and 0.232, respectively (47,48). The validity of the obtained complex structure was assessed using MolProbity (49). Data collection details and refinement statistics are provided in Table S3. All structural figures were prepared using PyMOL (Schrödinger, LLC).

### Database search

To search for the identified sequences in human collagen, we used these sequences as queries in the UniProt database (https://www.uniprot.org/blast).

## Supporting information

data set

## Data, materials, and software availability

Data of the structure of the human integrin a2I domain in the complex with GFOG**M**Rshort collagen peptide have been deposited in the Protein Data Bank (https://www.rcsb.org) (Accession code 9VB8).

## Acknowledgments

This work was supported by JST SPRING, Grant Number JPMJSP2128. We thank Edanz (https://jp.edanz.com/ac) for editing a draft of this manuscript.

## Notes

### Competing Interest Statement

The authors have declared no competing interest.

### Summary of Updates

The style of reference list has been updated.

