## Supporting Information for "Triple-helical ligands selectively targeting the closed αI domain of integrin α2β1"

Table S1. Theoretical occurrence rate of amino acid residues in the peptide library derived from NNK codon.

| Amino acid | ratio |
| --- | --- |
| Ala  Cys  Asp  Glu  Phe  Gly  His  Ile  Lys  Leu  Met  Asn  Pro  Gln  Arg  Ser  Thr  Val  Trp  Tyr  stop | 0.06250  0.03125  0.03125  0.03125  0.03125  0.06250  0.03125  0.03125  0.03125  0.09375  0.03125  0.03125  0.06250  0.03125  0.09375  0.09375  0.06250  0.06250  0.03125  0.03125  0.03125 |

Table S2. Amino acid sequences of synthesized peptides.

| Name | Sequence |
| --- | --- |
| C3-GFOG**M**R  C3-GFOGER  C3-PRG  Sol-GFOG**M**R  Sol-GFOG**M**R(^D^Pro)  Sol-GFOG**L**R  Sol-GFOG**I**R  GFOG**M**Rshort | H-Cys-Cys-Cys-(Pro-Hyp-Gly)_3_-Pro-Hyp-Gly-Phe-Hyp-Gly-Met-Arg-Gly-(Pro-Hyp-Gly)_4_-Cys-Cys-Cys-OH  H-Cys-Cys-Cys-(Pro-Hyp-Gly)_3_-Pro-Hyp-Gly-Phe-Hyp-Gly-Glu-Arg-Gly-(Pro-Hyp-Gly)_4_-Cys-Cys-Cys-OH  H-Cys-Cys-Cys-(Pro-Hyp-Gly)_5_-Pro-Arg-Gly-(Pro-Hyp-Gly)_4_-Cys-Cys-Cys-OH  H-(Pro-Hyp-Gly)_4_-Pro-Hyp-Gly-Phe-Hyp-Gly-Met-Arg-Gly-(Pro-Hyp-Gly)_5_-Pro-Tyr-NH_2_  H-(Pro-Hyp-Gly)_2_-^D^Pro-Hyp-Gly-Pro-Hyp-Gly-Pro-Hyp-Gly-Phe-Hyp-Gly-Met-Arg-Gly-(Pro-Hyp-Gly)_2_-^D^Pro-Hyp-Gly-(Pro-Hyp-Gly)_2_-Pro-Tyr-NH_2_  H-(Pro-Hyp-Gly)_4_-Pro-Hyp-Gly-Phe-Hyp-Gly-leu-Arg-Gly-(Pro-Hyp-Gly)_5_-Pro-Tyr-NH_2_  H-(Pro-Hyp-Gly)_4_-Pro-Hyp-Gly-Phe-Hyp-Gly-Ile-Arg-Gly-(Pro-Hyp-Gly)_5_-Pro-Tyr-NH_2_  Ac-(Gly-Pro-Hyp)_3_-Gly-Phe-Hyp-Gly-Met-Arg-(Gly-Pro-Hyp)_3_-NH_2_ |

(^D^Pro indicates D-proline.)

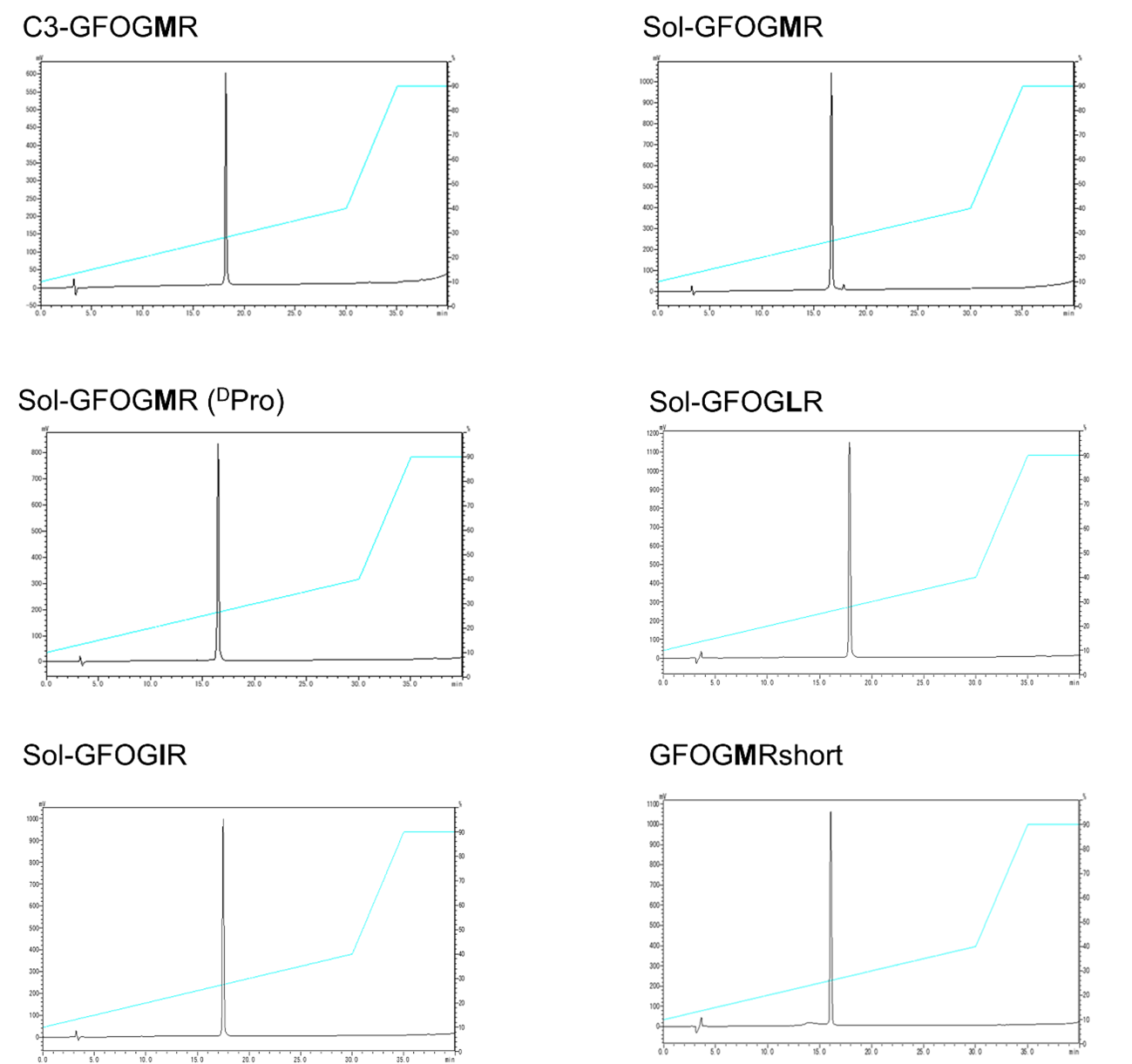

Figure S1. Reverse phase-high performance liquid chromatography (RP-HPLC) profiles of synthesized peptides. Column: Cosmosil 5C18-ARII (4.6 i.d. × 250 mm). Gradient: 10%−40% CH_3_CN in 0.05% TFAaq. over 30 min at 60 ^o^C. Flow rate: 1.0 mL/min. Absorbance: 220 nm.

C3-GFOG**M**R

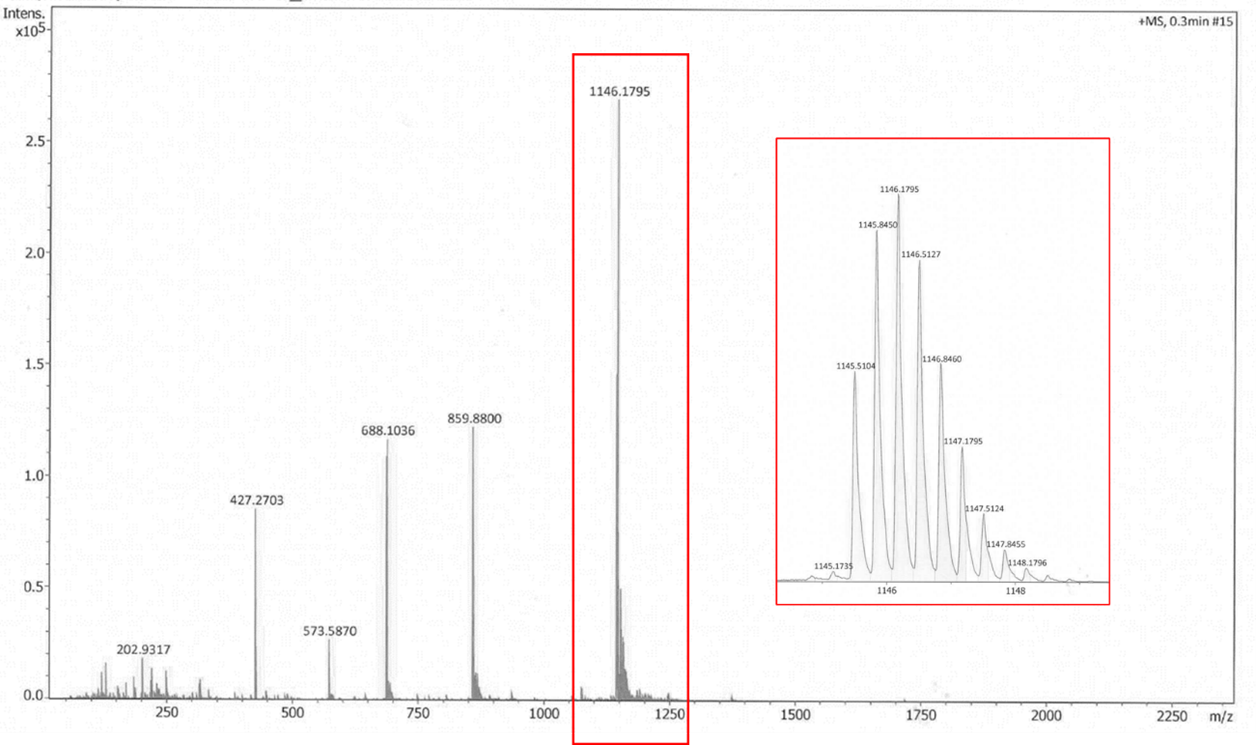

calcd. MS [M_m_ + 3H] ^3+^: 1146.659	found: 1146.180

Sol-GFOG**M**R

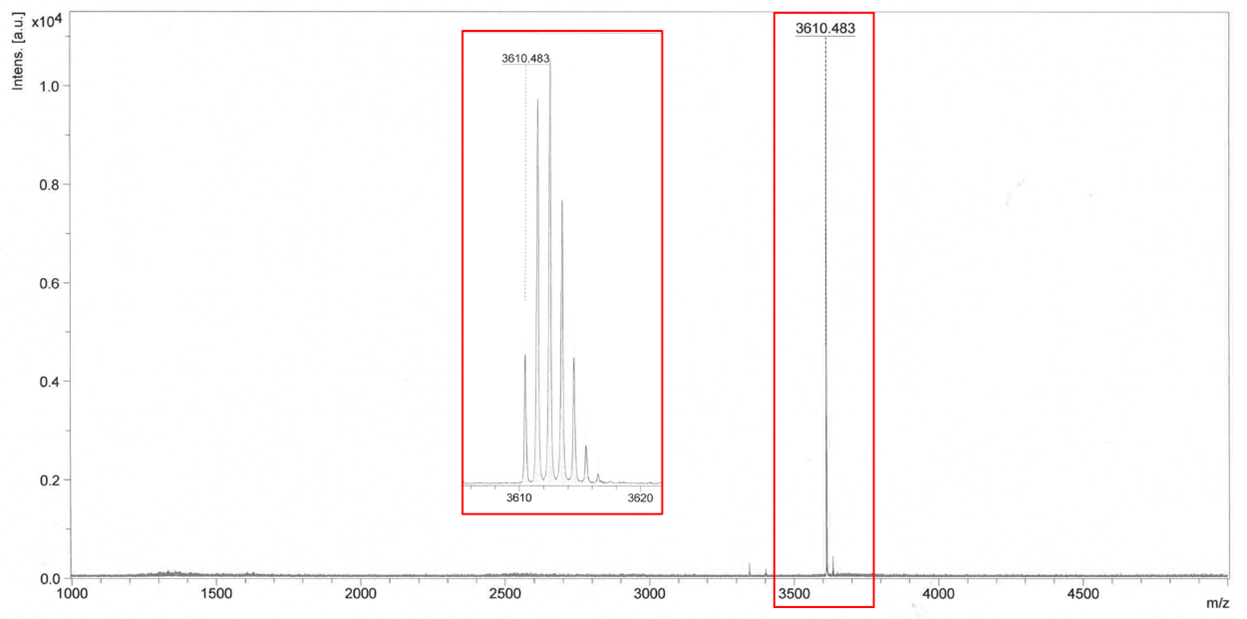

calcd. MS [M_m_ + H]^+^: 3610.675	found: 3610.483

Figure S2. Mass spectrometry charts of synthesized peptides.

Sol-GFOG**M**R(^D^Pro)

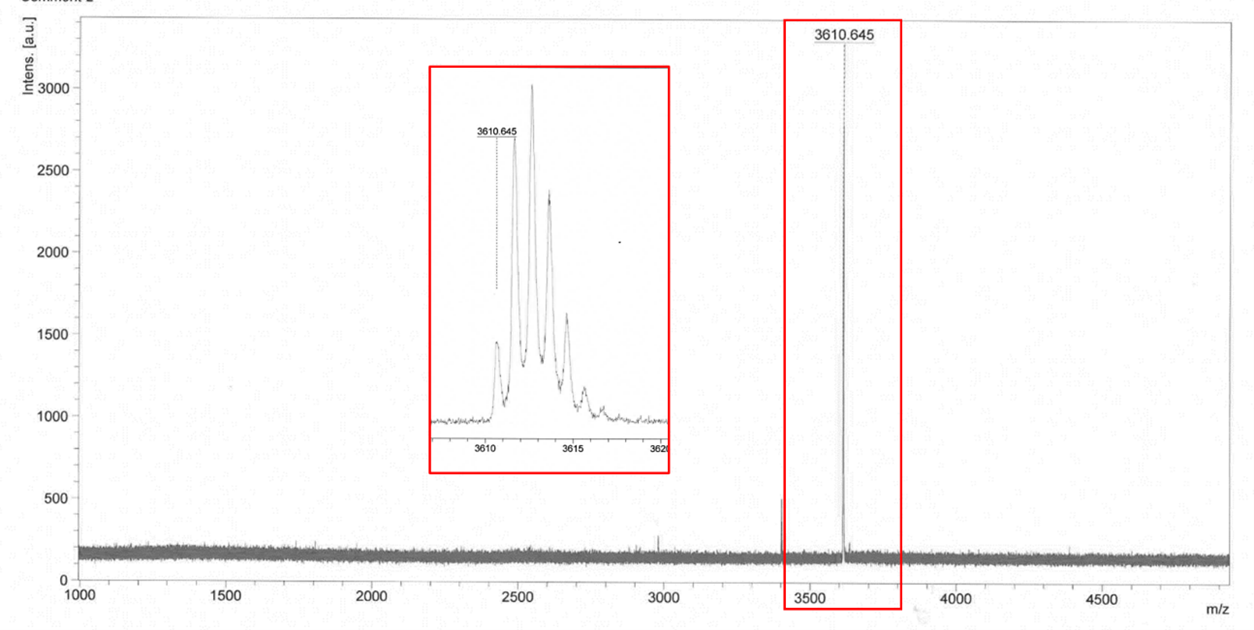

calcd. MS [M_m_ + H]^+^: 3610.675	found: 3610.645

Sol-GFOG**L**R

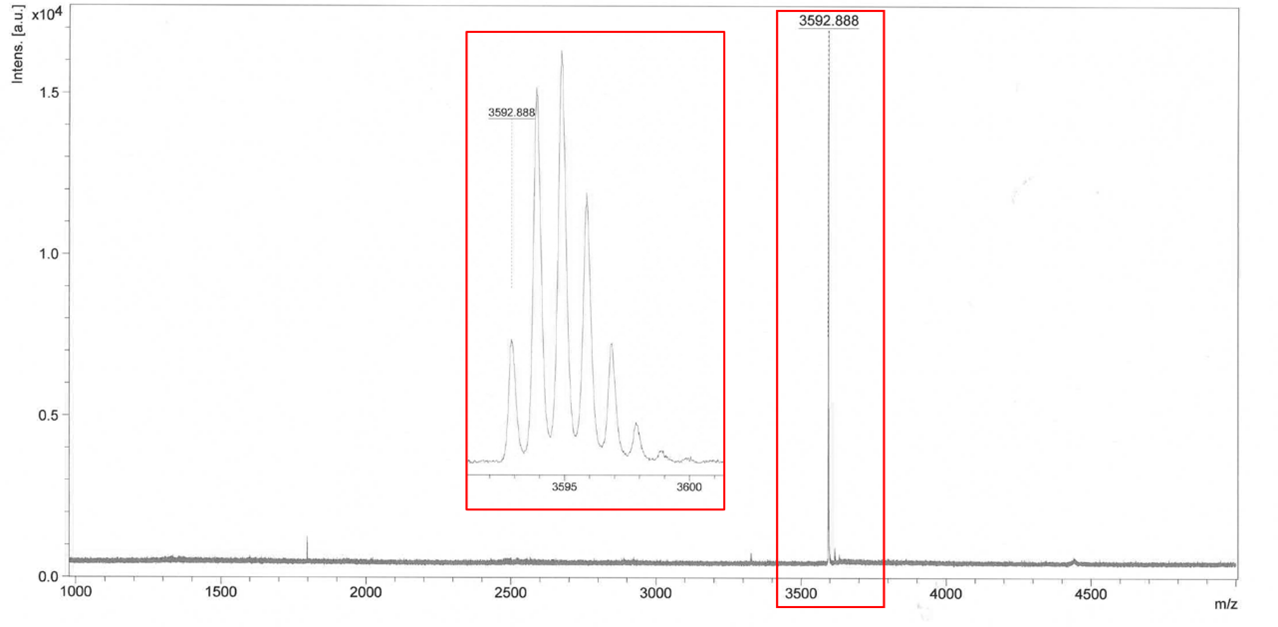

calcd. MS [M_m_ + H]^+^: 3592.713	found: 3592.888

Figure S2. (*Continued*)

Sol-GFOG**I**R

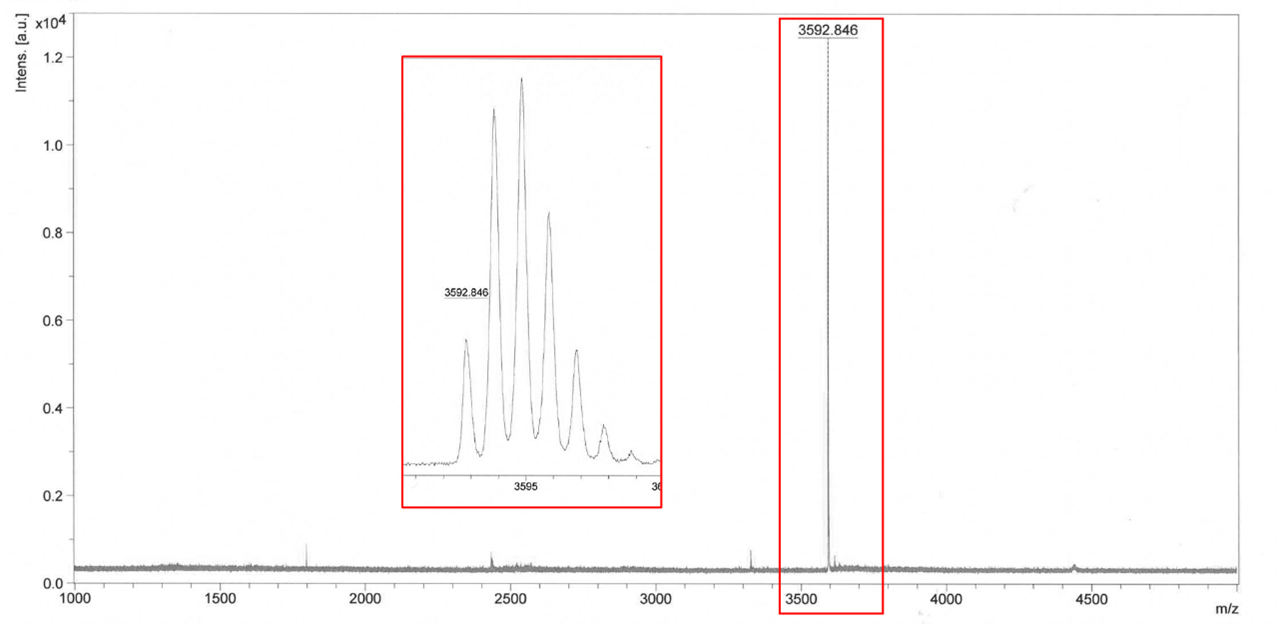

calcd. MS [M_m_ + H]^+^: 3592.713	found: 3592.846

GFOG**M**Rshort

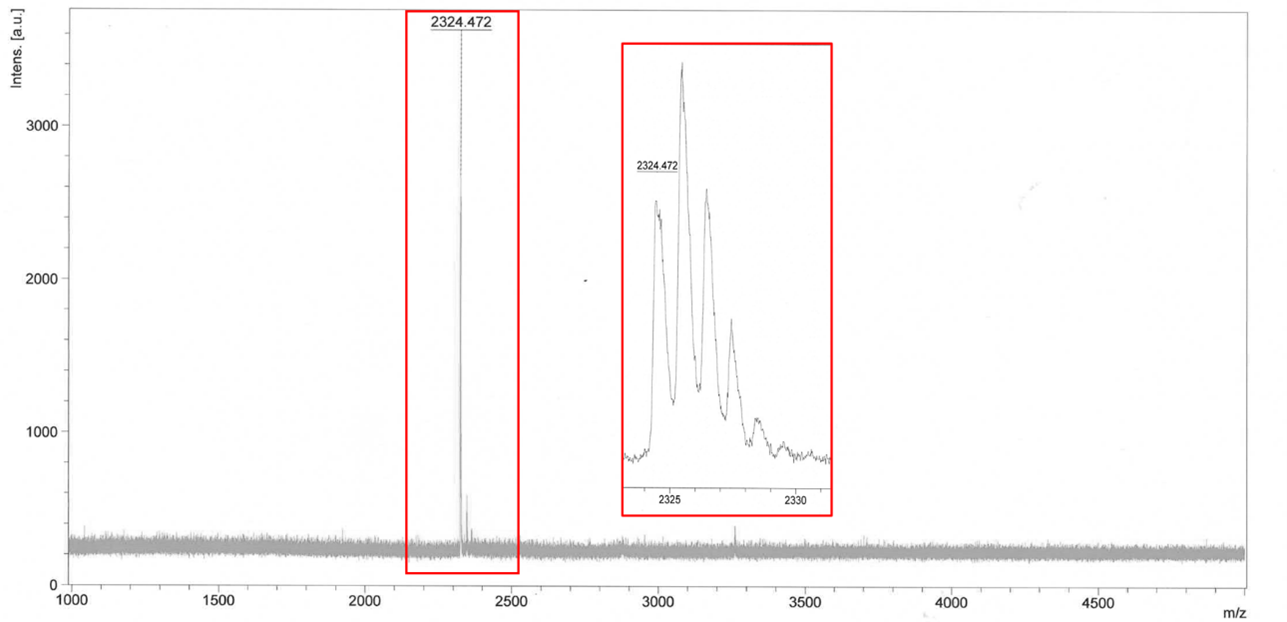

calcd. MS [M_m_ + H] ^+^: 2324.077	found: 2324.472

Figure S2. (*Continued*)

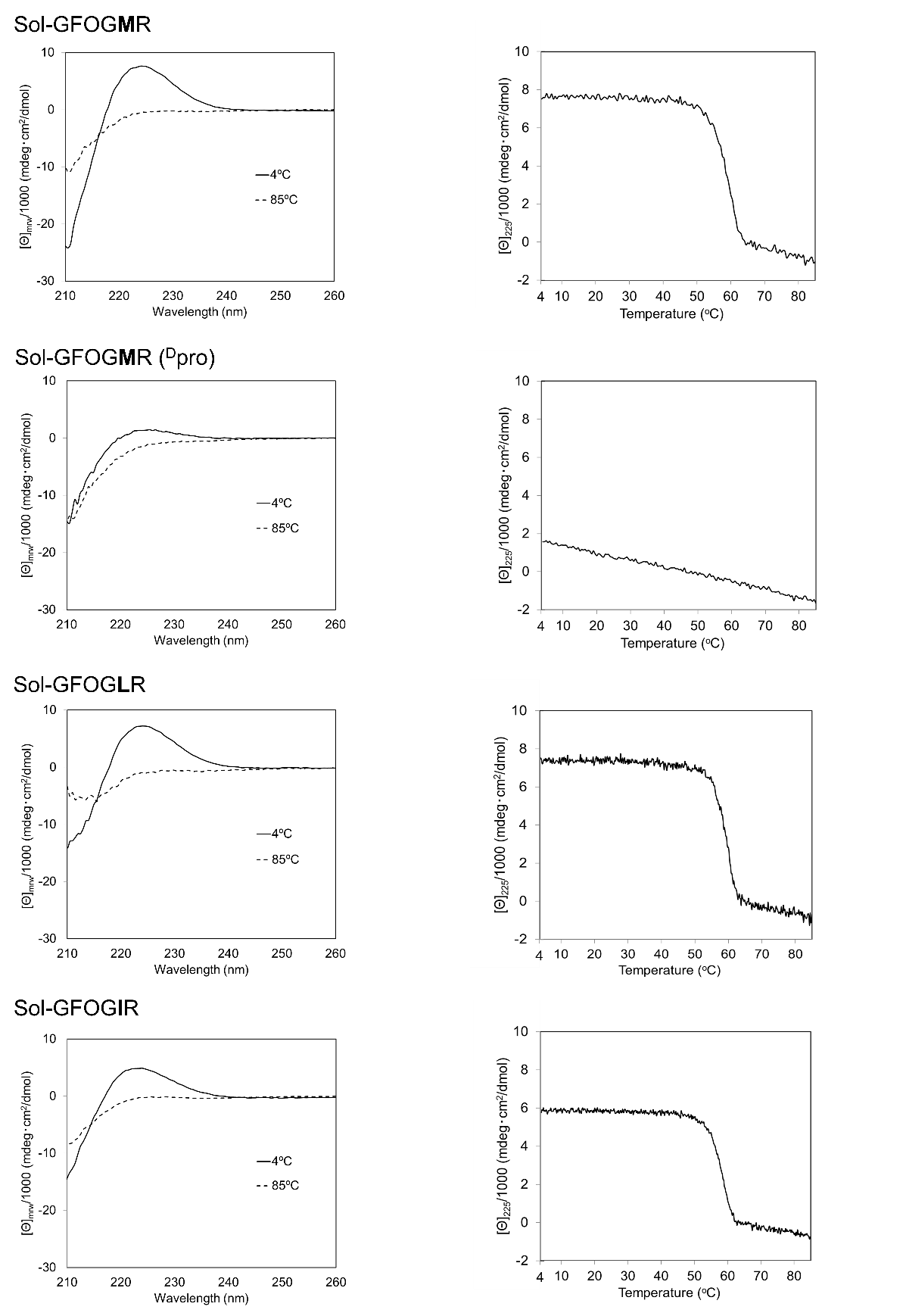

Figure S3. CD profiles of peptides at 4 ℃ (solid lines) and 85 ℃ (dotted line) (left panels). CD spectra recorded at 4 and 85 °C (Right panels). A positive maximum at 225 nm indicates a polyproline II (PP-II)-like helical structure (1). A cooperative decrease in the signal with increasing temperature indicates the melting of the triple helix. A linear decrease in signal indicates a single-chain polyproline II helix.

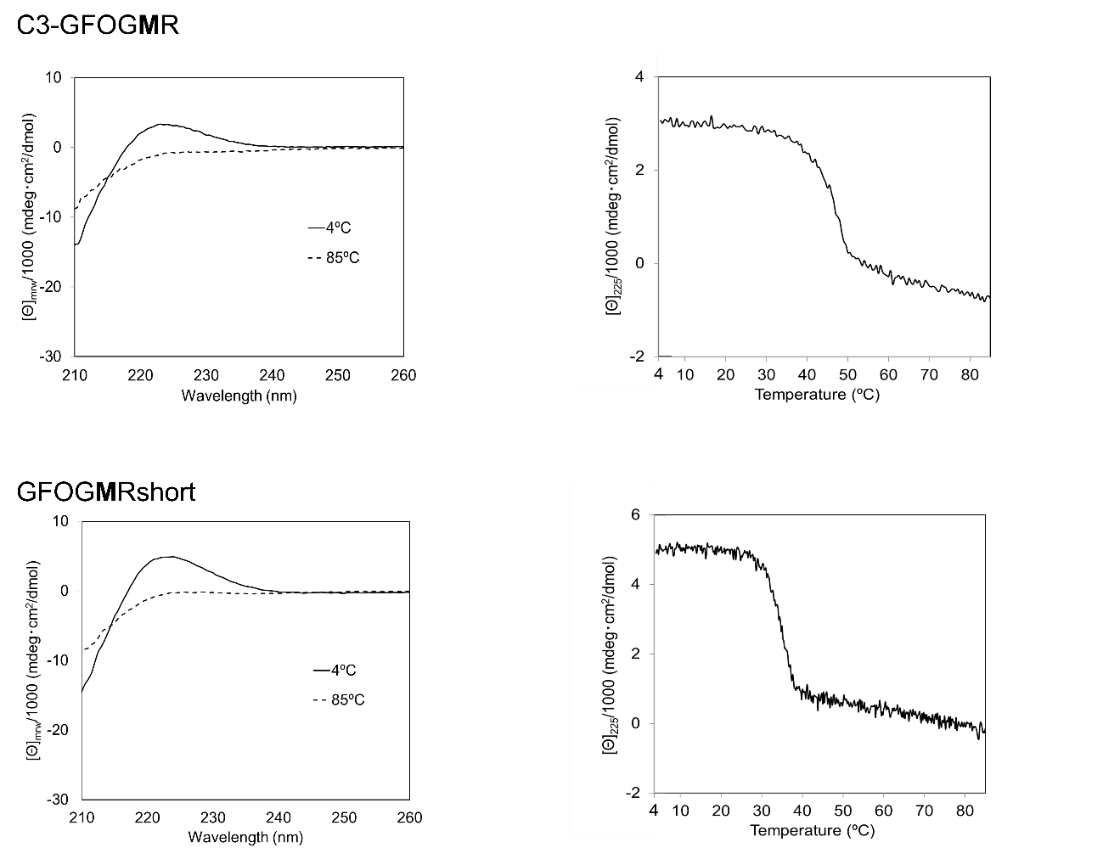

Figure S3. (*Continued*).

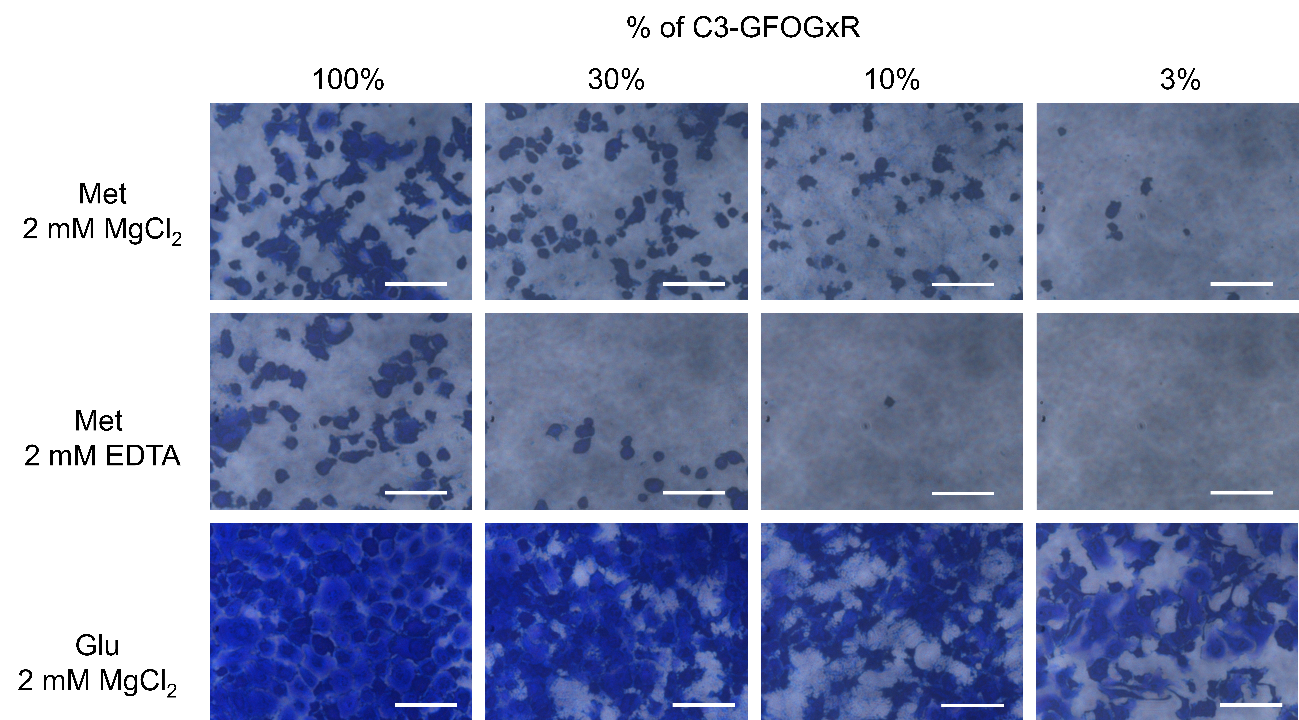

Figure S4. HT1080 cell adhesion to triple-helical peptide co-polymers in a sequence-dependent manner. Adhesion of C3-GFOGxR (x = Glu, Met) and C3-PRG co-polymer in the presence of 2 mM MgCl₂ or 2 mM EDTA. Adherent cells were stained with crystal violet and observed under a phase-contrast microscope. "% of C3-GFOGxR" represents the proportion of C3-GFOGxR in the total peptide weight. Scale bar represents 100 µm.

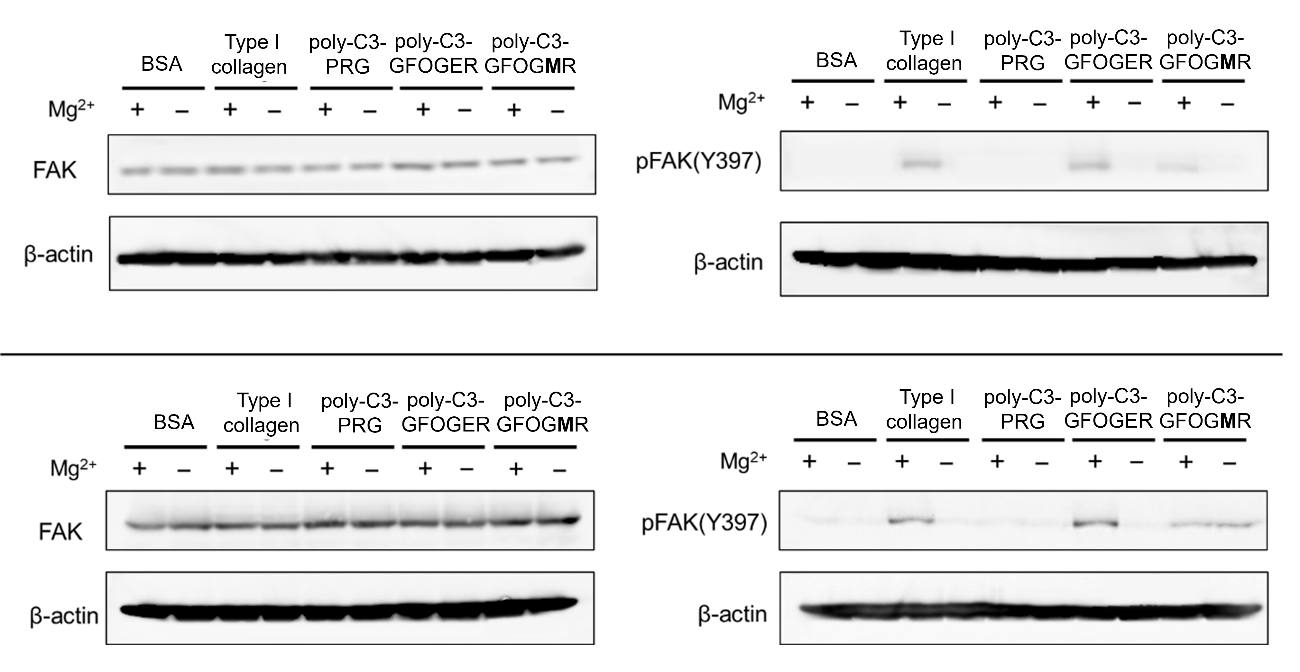

Figure S5. Western blotting of FAK and phosphorylated FAK (Y397). The phosphorylation level of FAK shown in Figure 4D was measured using the results from Figure 4C and the two additional results presented in this figure. Mg²⁺ + and − indicate that HT1080 cells were cultured in the presence of 2 mM MgCl₂ or 2 mM EDTA, respectively.

Table S3. Data collection and refinement statistics.

|  | Human integrin α2I domain-GFOG**M**Rshort complex |
| --- | --- |
| **Data collection**  Beamline  Detector  Wavelength (Å)  Space group  Unit-cell parameters  *a, b, c* (Å)  *α*, *β*, γ (º)  Unique reflections  Resolution (Å)  *R_mead_* (%)  CC_1/2_ (%)  Completeness (%)  Average *I/σ* (I)  Redundancy  **Refinement**  Resolution (Å)  Reflections  No. Atoms  Protein  Waters  *R_work_*  *R_free_*  R.M.S.D. from ideal  Bonds (Å)  Angles (º)  *B*-factors (Å^2^)  Ramachandran plot analysis (%)  Most favored  Allowed | SPring-8 BL45XU  PILATUS 6M  1.0  P1  60.36, 71.76, 83.86,  110.28, 90.09, 112.21  153,836 (7,439)  48.67-1.60 (1.63-1.60)  15.3 (70.8)  97.8 (50.8)  96.5 (94.1)  4.7 (1.5)  2.3 (2.3)  48.67-1.60 (1.62-1.60)  153819 (5027)  8233  1505  0.185 (0.324)  0.222 (0.361)  0.008  1.061  25.19  98.78  1.22 |

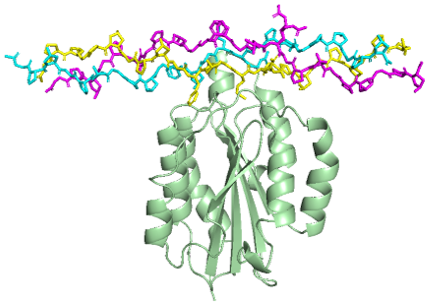

Figure S6. The α2I domain-GFOG**M**Rshort complex is shown with the α2I domain as a cartoon and GFOG**M**Rshort as a stick model. Leading (L), middle (M), and trailing(T) chains in the triple helix are colored cyan, yellow, and magenta, respectively.

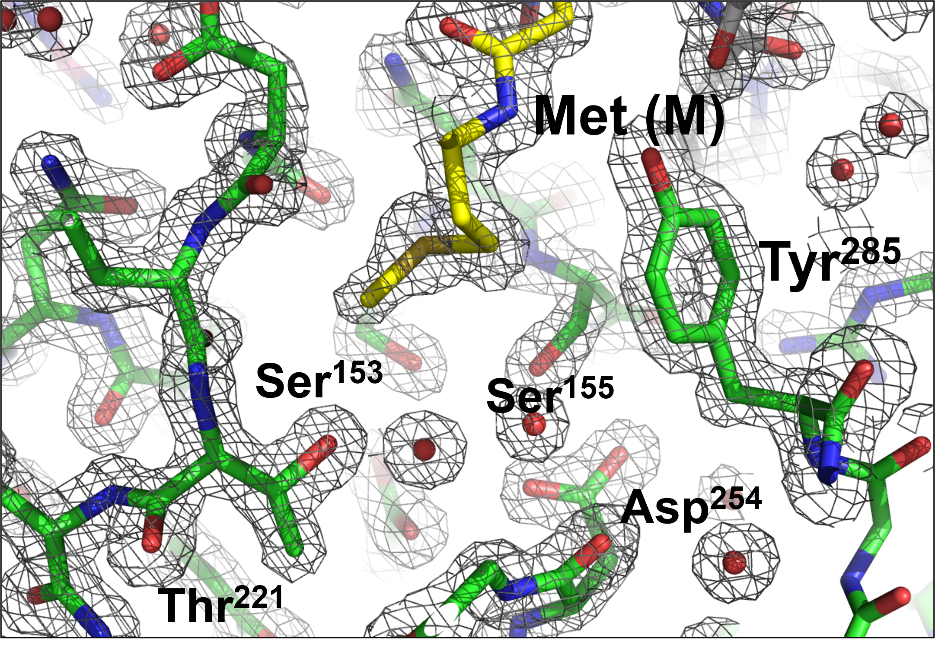

Figure S7. 2mFo-DFc electron density map (gray mesh) around the MIDAS motif in the complex of GFOG**M**Rshort and the α2I domain, contoured at 2.0 σ.

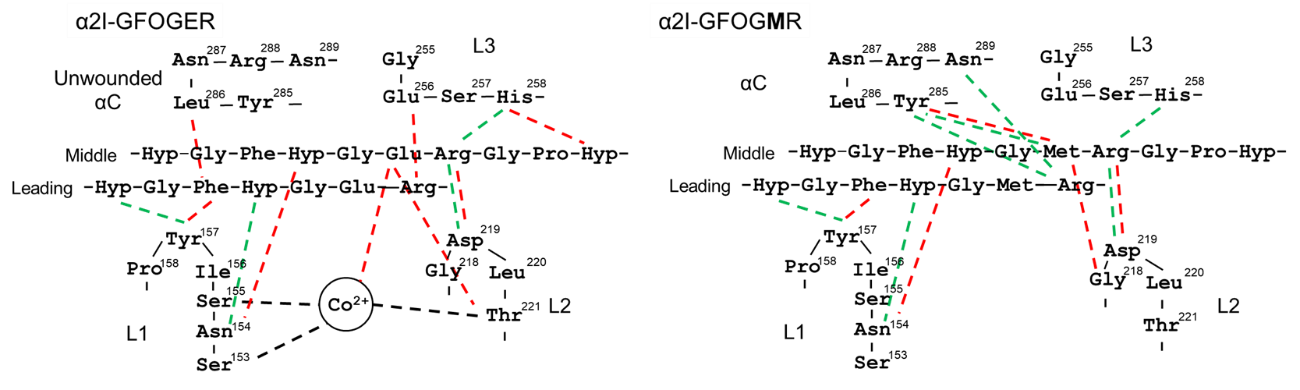

Figure S8. Schematic illustration of the interactions between peptides and the α2I domain. Interactions of the backbone of the triple-helical peptide are shown in green dotted lines, and interactions of the side chains are shown in red lines.

Table S4. Binding sequences for the α2I T221A mutant obtained from Y2H screening.

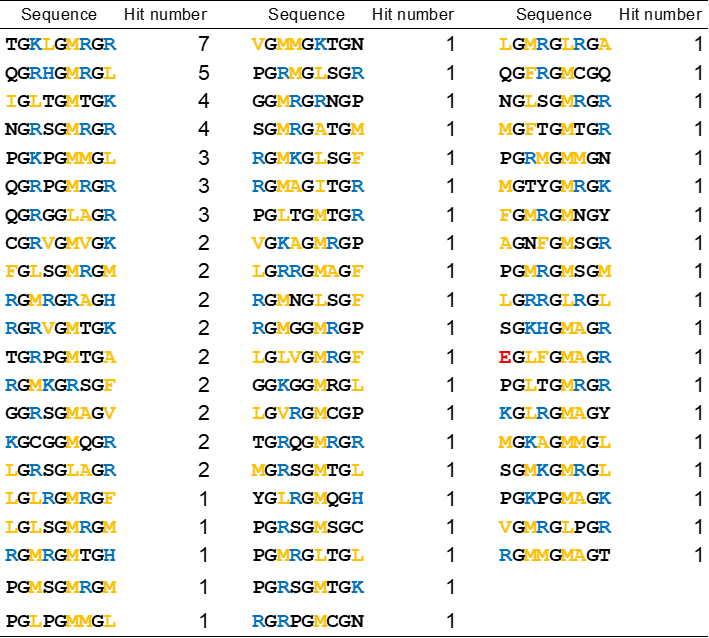

Table S5. Binding sequences for the α2I E318W mutant obtained from Y2H screening.

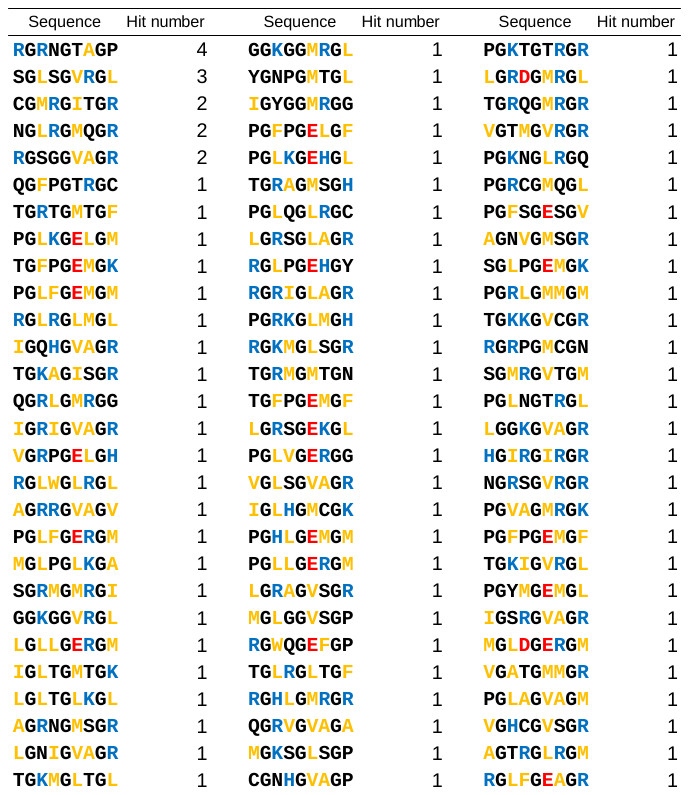

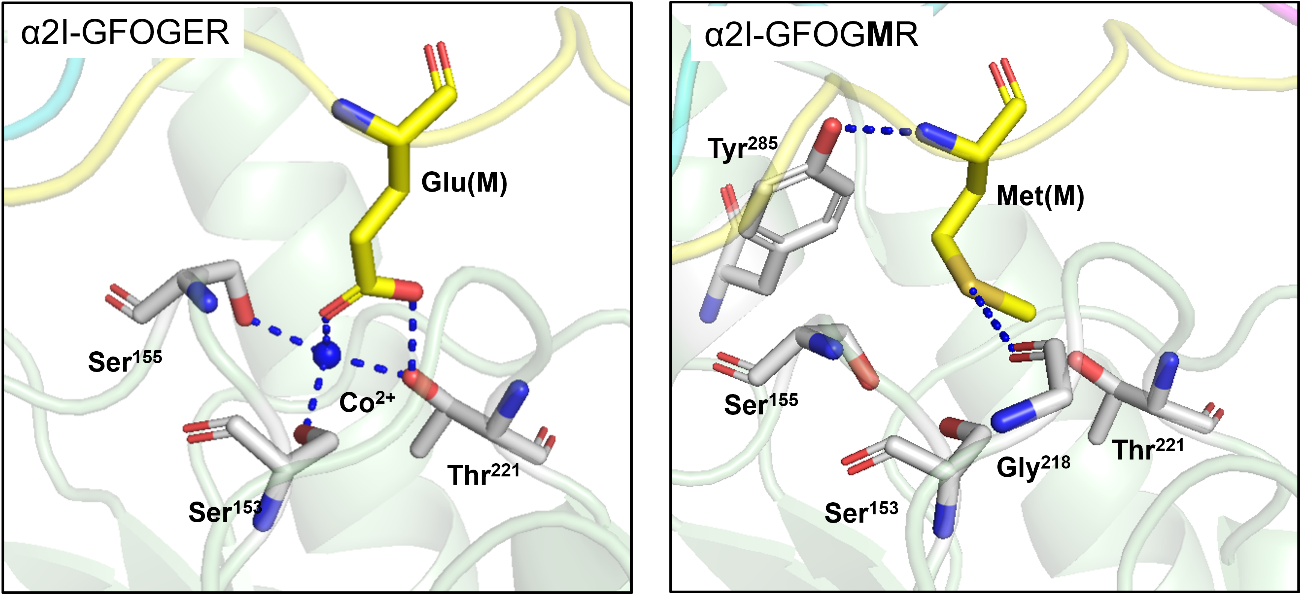

Figure S9. Close-up views of the MIDAS in the α2I-peptide complexes. The right panel shows the complex of the α2I domain with GFOG**M**Rshort, while the left panel shows the complex of the α2I domain with a GFOGER-containing peptide (PDB ID code: 1DZI). Leading (L), middle (M), and trailing (T) chains in the triple helix are colored cyan, yellow, and magenta, respectively.

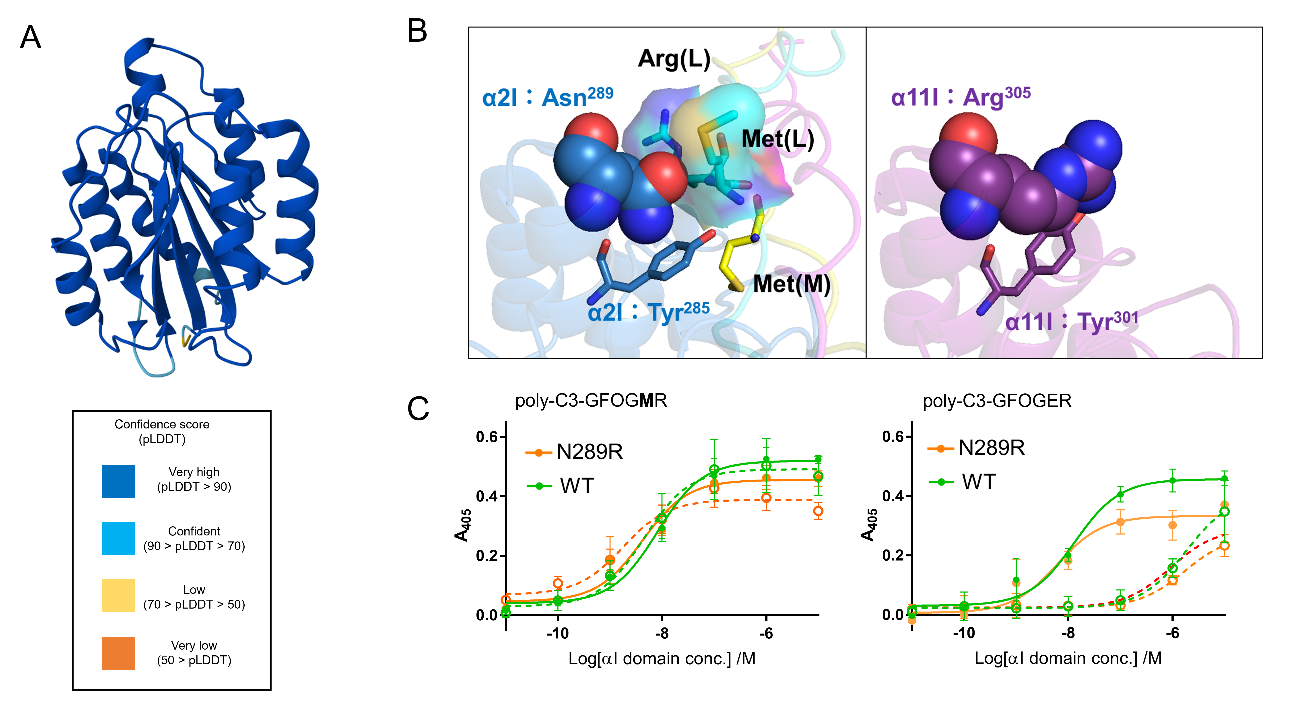

Figure S10. Structural comparison between integrin α2I and α11I domain. (A) Structures of integrin α11I domain predicted by AlphaFold3 (colored by pLDDT score) (2). The predicted α11I domain adopts a closed conformation. (B) Comparison of regions corresponding to Tyr285 and Asn289 of the α2I domain in α11I domain. The α2I domain (blue, left panel) is in the complex with GFOG**M**Rshort. The structures of the α11I (purple, right panel) domains are predicted using AlphaFold3. Leading (L), middle (M), and trailing (T) chains in the triple helix are colored cyan, yellow, and magenta, respectively. Asn289 in the α2I domain and the corresponding Arg305 in the α11I domain are shown as spheres. The Arg and Met residues in the leading chain of GFOG**M**Rshort that contact Asn289 are shown the surfaces. (C) Interaction with surface immobilized C3-peptide polymer. Binding of N289R (yellow line) and WT (green line). Adhesion was conducted in the presence of either 2 mM MgCl2 (solid line) or 5 mM EDTA (dotted line). n = 3, mean ± S.D.

Table S6. Sequences of primers for preparing α2I domain mutants.

| Name | Sequence |
| --- | --- |
| α2I T221A forward  α2I T221A reverse  α2I E318W forward  α2I E318W reverse  α2I Y285F forward  α2I Y285F reverse  α2I Y285A forward  α2I Y285A reverse  α2I N289R forward  α2I N289R reverse | 5′-GATTTGGCGAATACTTTTGGTGCTATT-3′  5′-AGTATTCGCCAAATCACCGCCATATTG-3′  5′-TCAGATTGGGCTGCGTTATTGGAGAAA-3′  5′-CGCAGCCCAATCTGAGACATTGAAGAA-3′  5′-CTTGGCTTTTTAAACCGTAATGCCCTA-3′  5′-GTTTAAAAAGCCAAGAACTGCAATTCC-3′  5′-CTTGGCGCTTTAAACCGTAATGCCCTA-3′  5′-GTTTAAAGCGCCAAGAACTGCAATTCC-3′  5′-AACCGTAGAGCCCTAGATACCAAGAAC-3′  5′-TAGGGCTCTACGGTTTAAGTAGCCAAG-3′ |

Table S7. Sequences of primers for preparing α2I domain for crystallization.

| Name | Sequence |
| --- | --- |
| α2I cryst N forward  α2I cryst N reverse  α2I cryst C forward  α2I cryst C reverse | 5′-phospho-AGCCCATCCTTAATTGACGTCG-3′  5′-ACGACCTTCGATCAGATCC-3′  5′-ATCGTGACTGACCTGACGATC-3′  5′-phosho-TTAACCTTCAATGGAAAAGATCTGTTC-3′ |

Table S8. Sequences of ssDNA for preparing prey constructions in Figure 1B.

| Name | Sequence |
| --- | --- |
| Seq. 1 forward  Seq. 1 reverse  Seq. 2 forward  Seq. 2 reverse  Seq. 3 forward  Seq. 3 reverse  Seq. 4 forward  Seq. 4 reverse  Seq. 5 forward  Seq. 5 reverse  Seq. *a* forward  Seq. *a* reverse | 5′-CCCAGGTTTTCCAGGTGAAAGAGGTCCTC-3′  5′-CCGGGAGGACCTCTTTCACCTGGAAAACCTGGGGGCC -3′  5′-CCAAGGTTTCGTTGGTATGTCTGGTAGAC-3′  5′-CCGGGTCTACCAGACATACCAACGAAACCTTGGGGCC -3′  5′-CAGAGGTAAGTTCGGTATGAGAGGTAGAC -3′  5′-CCGGGTCTACCTCTCATACCGAACTTACCTCTGGGCC -3′  5′-CGCTGGTTTGATTGGTTTGAGAGGTCATC -3′  5′-CCGGGATGACCTCTCAAACCAATCAAACCAGCGGGCC -3′  5′-CAGAGGTGTTGGTGGTATGAAGGGTAGAC -3′  5′-CCGGGTCTACCCTTCATACCACCAACACCTCTGGGCC -3′  5′-CATGGGTTTCCCAGGTATGAGAGGTACTC -3′  5′-CCGGGAGTACCTCTCATACCTGGGAAACCCATGGGCC -3′ |

Table S9. Sequences of primers for Sanger sequencing.

| Name | Sequence |
| --- | --- |
| pgad seq primer  3ad rv | 5′-CCACTACAATGGATGATGTATATAA -3′  5′-AGATGGTGCACGATGCACAG-3′ |

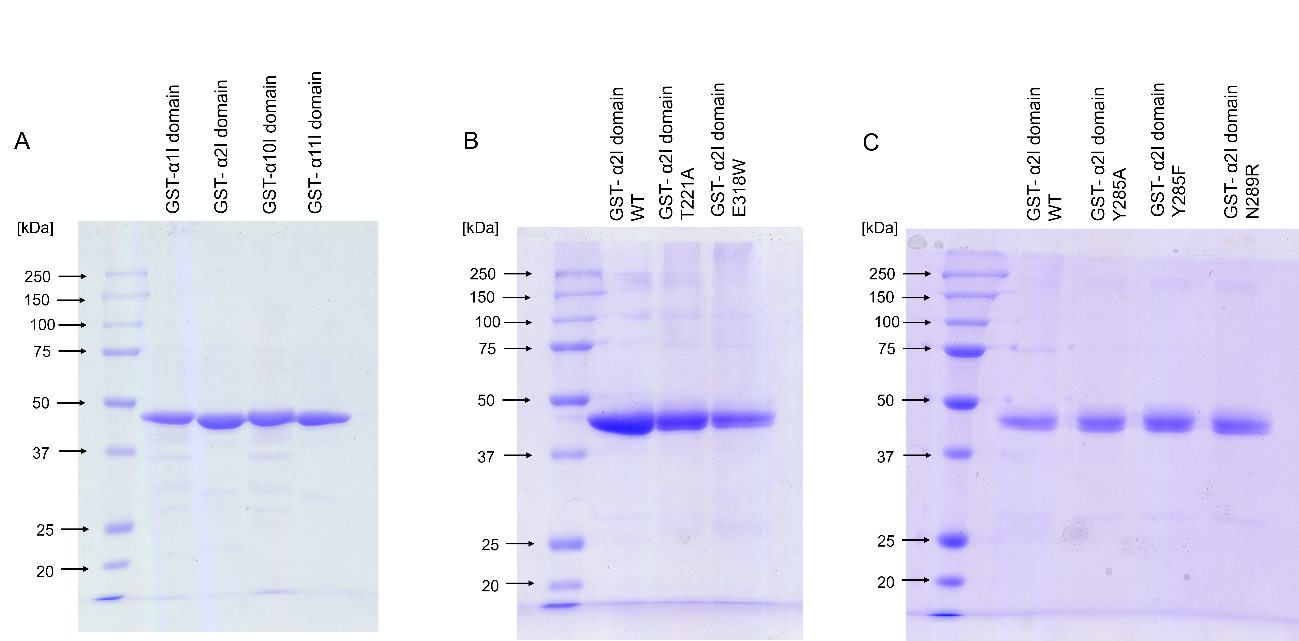

Figure S11. SDS-PAGE analysis of purified GST-αI domain. Proteins underwent electrophoresis with 12% acrylamide gel, and bands were visualized by Coomassie brilliant blue staining. The molecular weights are shown in kilodalton (kDa). The molecular weights of GST-α1, α2, α10, and α11 I domains were 50, 48, 49, and 49 kDa, respectively.
